# Fluoxetine targets an allosteric site in the enterovirus 2C AAA+ ATPase and stabilizes the hexameric complex

**DOI:** 10.1101/2021.04.26.440876

**Authors:** Daniel L. Hurdiss, Priscila El Kazzi, Lisa Bauer, Nicolas Papageorgiou, François P. Ferron, Tim Donselaar, Arno L.W. van Vliet, Bruno Canard, Etienne Decroly, Andrea Brancale, Tzviya Zeev-Ben-Mordehai, Friedrich Förster, Frank J.M van Kuppeveld, Bruno Coutard

**Affiliations:** Virology Section, Infectious Diseases and Immunology Division, Department of Biomolecular Health Sciences, Faculty of Veterinary Medicine, Utrecht University, 3584CL Utrecht, The Netherlands; Cryo-Electron Microscopy, Bijvoet Center for Biomolecular Research, Department of Chemistry, Faculty of Science, Utrecht University, Padualaan 8, 3584 CH Utrecht, The Netherlands; Aix Marseille Université, CNRS, AFMB UMR 7257, Marseille, France; School of Pharmacy & Pharmaceutical Sciences, Cardiff University, King Edward VII Avenue, Cardiff CF10 3NB, United Kingdom; Unité des Virus Émergents (UVE: Aix-Marseille Univ-IRD 190-Inserm 1207), Marseille, France

## Abstract

The enterovirus genus encompasses many clinically important human pathogens such as poliovirus, coxsackieviruses, echoviruses, numbered enteroviruses and rhinoviruses. These viruses are the etiological agents of several human diseases, including hand-foot-and-mouth disease, neonatal sepsis, encephalitis, meningitis, paralysis and respiratory infections. There is an unmet need for antivirals to treat these diseases. The non-structural protein 2C is a AAA+ helicase and plays a key role in viral replication. As such, it is an attractive target for antiviral drug development. Several repurposing screens with FDA-approved drugs have identified 2C-targeting compounds such as fluoxetine and dibucaine, but the molecular basis of 2C inhibition has remained enigmatic. Here we present the 1.5 Å resolution crystal structure of the soluble fragment of coxsackievirus B3 2C protein in complex with (S)-fluoxetine (SFX), which reveals a conserved, hydrophobic drug-binding pocket which is distal to the ATP binding site. To decipher the molecular mechanism of inhibition by fluoxetine and other 2C-targeting compounds, we engineered a soluble, hexameric and ATPase competent 2C protein. Using this system, we show that SFX, dibucaine, HBB and guanidine hydrochloride inhibit 2C ATPase activity in a dose-dependent manner. Moreover, using cryo-EM analysis, we demonstrate that SFX and dibucaine lock 2C in a defined hexameric state, rationalizing their mode of inhibition and allowing us to generate the first reconstruction of the oligomeric complex. Taken together, these results provide important structural and mechanistic insights into 2C inhibition and provide a robust engineering strategy which can be used for structural, functional and drug-screening analysis of 2C proteins from current or future enteroviruses.

## Introduction

Viruses belonging to the genus *Enterovirus*, within the family *Picornaviridae*, are responsible for a broad range of diseases. This genus contains four clinically relevant enterovirus (EV) species (EV-A-D) and three rhinovirus species (RV-A-C). Notable members of these species include polioviruses, coxsackieviruses, echoviruses and numbered EVs (e.g. EV-A71 and EV-D68). EV-associated diseases include respiratory infections, hand-foot-and-mouth disease, conjunctivitis, viral myocarditis, pancreatitis, aseptic meningitis, encephalitis, and acute flaccid paralysis^1^. RVs cause the common cold but can also trigger the exacerbation of asthma and chronic obstructive pulmonary disease^2^. While most EV infections are mild and self-limiting, fatal complications may arise in immunocompromised patients and young children^3^. Outbreaks of EV-A71 and EV-D68 represent major public health concerns because these viruses are associated with severe neurological complications^4–6^. Vaccines are only available for poliovirus and EV-A71, with the latter only approved in China^7,8^. Treatment of other clinically important EVs is currently limited to supportive care. Given the large numbers of serotypes (>100 EVs and >200 RVs), the development of a pan-EV and RV vaccine seems unfeasible. This underscores the urgent clinical need for the development of broad-spectrum antivirals to treat EV-associated diseases.

The EV genome encodes seven non-structural proteins (2A-2C and 3A-3D), which are involved in replication, and four structural proteins (VP1-VP4) which form the capsid. Several direct acting agents (DAA) targeting either the capsid, the viral polymerase (3D^pol^) or the viral protease (3C^pro^) have been identified. While some of these compounds reached clinical trials, their development was stopped due to limited efficacy, poor bioavailability or toxicity issues^9^. An alternative target for antiviral drug discovery is the non-structural protein 2C which exhibits several important hallmarks of a promising DAA target. 2C is functionally indispensable and plays an essential role in several steps in the EV life cycle, e.g. viral RNA replication and encapsidation^10^. Moreover, 2C is highly conserved^11^, raising the possibility for broad-spectrum DAA development.

2C belongs to the helicase superfamily 3 (SF3) and acts as an ATP-dependent RNA helicase and ATP-independent RNA remodeler^12^. The role of this enzyme in viral replication likely requires an oligomeric organization, which has already been observed for recombinant 2C proteins^13–15^. Previous studies suggested that the functional oligomerization state of 2C is hexameric, consistent with other members of the SF3 helicase family^16,17^. Structural and functional characterization of the full-length oligomeric 2C protein remains challenging due to the N-terminal amphipathic helix which renders the protein poorly soluble^13–15,18,19^. Recent structures of a soluble, monomeric fragment of 2C from EV-A71 and PV provided molecular details of the adenosine triphosphatase (ATPase) domain, a cysteine-rich zinc finger and a carboxyl-terminal helical domain^16,17^. Nevertheless, a structure of the oligomeric 2C remains elusive.

Given the central role of 2C during viral replication, it is not surprising that several compounds targeting 2C, such as guanidine hydrochloride (GuaHCl), HBB, MRL-1237 and TBZE-029, have been identified^18,20–22^. Additionally, several drug-repurposing screens have uncovered FDA-approved drugs such as fluoxetine, dibucaine, pirlindole and zuclopenthixol as inhibitors of EV-B and EV-D species, but their mode of action is not yet understood^23,24^. Fluoxetine (Prozac^®^) is a selective serotonin-reuptake inhibitor that is used clinically for the treatment of major depression and anxiety disorders^25^. Mode-of-action studies revealed that only the *S*-enantiomer of fluoxetine potently binds 2C and inhibits EV-B and D replication^26,27^. Despite the growing number of identified 2C-targeting compounds, the molecular basis for their antiviral activity remains unknown^28,29^. Such information would provide an invaluable resource for structure-based design of potent, pan-enterovirus antivirals with low toxicity.

Here, we present the 1.5 Å resolution crystal structure of the soluble fragment of CV-B3 2C in complex with SFX. The structure reveals a highly conserved hydrophobic pocket, distal to the ATP binding site, into which the SFX trifluoro-phenoxy moiety inserts. To functionally validate our structure and study the inhibitory mechanism of SFX and other 2C targeting compounds, we engineered a CV-B3 2C protein fused to a heterologous hexamerization domain. The chimeric protein recovered ATPase activity, allowing us to investigate the inhibitory effect of several antivirals targeting EV 2C, each of which displayed a dose-dependent ATPase inhibition. Moreover, incubation of our engineered 2C protein with SFX or dibucaine demonstrated that these drugs stabilize the hexameric complex. This allowed us to capture the first three-dimensional structure of the 2C hexamer by cryo-EM and suggests an inhibitory mechanism for these drugs. Taken together, these data provide new mechanistic insights into the mode-of-action of 2C targeting compounds and offer unique tools for the design and validation of 2C inhibitors.

## Results

### Crystal structure of CV-B3 Δ116-2C in complex with SFX

Inspired by recent structural reports of 2C protein fragments from EV-A and EV-C species^16,17^, we attempted a similar truncation strategy to obtain crystals of a fluoxetine-sensitive EV-B species member, namely CV-B3. Thermal denaturation experiments showed that SFX increased the melting temperatures (Tm + 1.51°C) of the CV-B3 Δ116-2C, confirming that this truncated protein can be bound by the drug (Figure S1). To understand how SFX binds to 2C, we attempted to crystallize our truncated 2C construct in the presence or absence of the drug. While we were able to obtain a high resolution (1.5 Å) structure of the complex (Table S1), no crystals were obtained for the apo 2C protein. This suggests that SFX stabilizes the 2C protein in a conformation prone to crystallization.

The overall fold of CV-B3 Δ116-2C is similar to the analogous structure from EV-A71 and poliovirus, comprising a Rossmann fold core domain, a zinc finger domain and a C-terminal helix (Figure 1A and S2). The Walker A motif is located on the loop connecting β1 and α1, forming a putative phosphate binding loop (P-loop). The Walker B motif is found between β3 and α2 while the Walker C motif, containing N223, is found at the tip of β4^30^. Two large flexible loops located between strands β2-β3 and α2-β4 shield one side of the central sheet. Together, these loops form a hydrophobic cavity into which the SFX molecule binds (Figure 1A-B and S2). The interaction between SFX and the CV-B3 2C is mainly mediated by hydrophobic interactions involving the side chains of residues L157, P159, M175, D176, L178, C179, P182 and D186 (Figure 1B-C). The complex is also stabilized by a hydrogen bond between the amide group of the C179 main chain and the hydroxyl group of SFX. Finally, in one of our structures, SFX interacts via its amino group with the main chain of D186 (Figure 1C).

**Figure 1.**
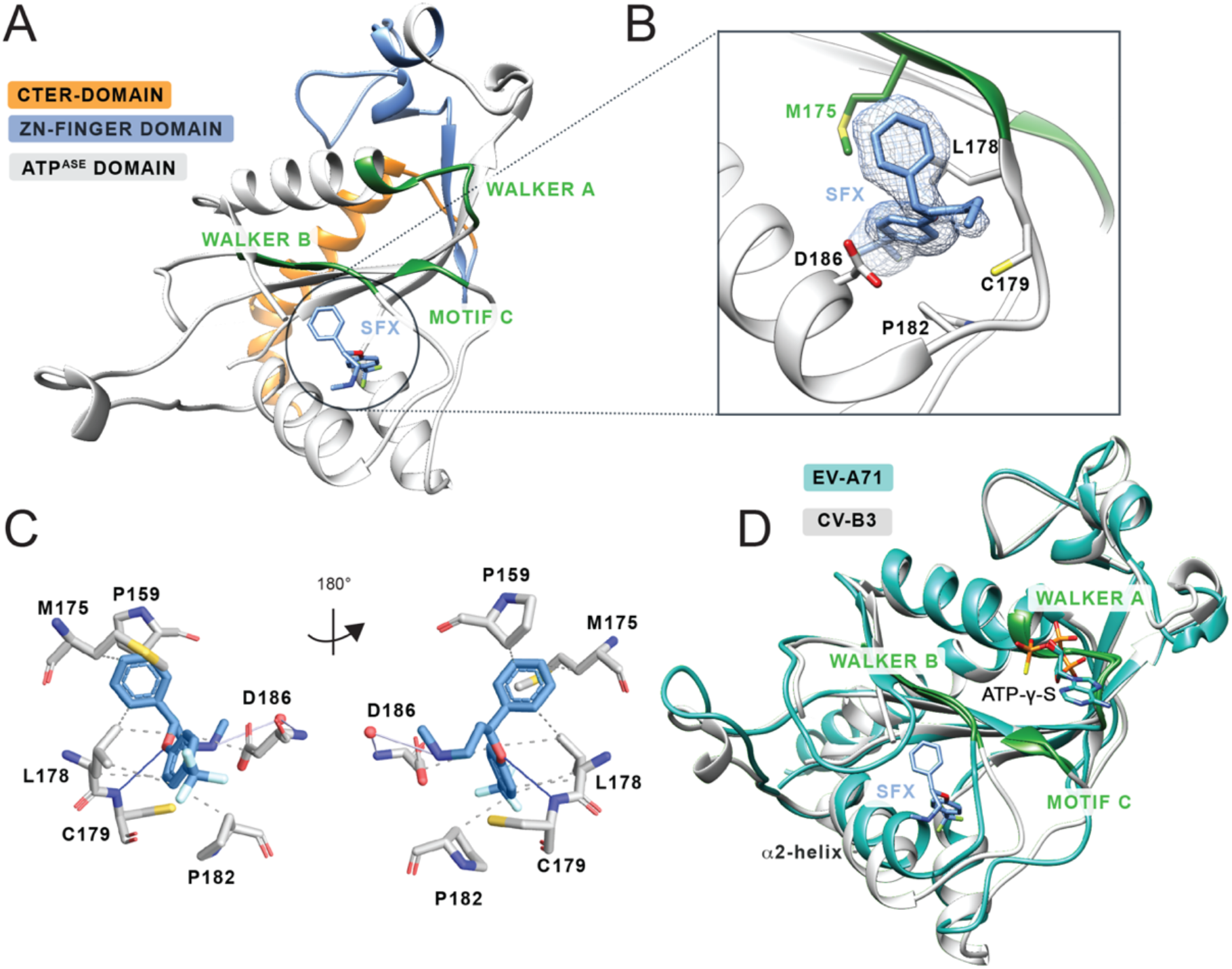
Crystal structure of CV-B3 Δ116-2C in complex with SFX. (A) The C-terminal part of 2C is highlighted in orange. The Zinc-Finger domain is highlighted in blue, and the ATPase domain is highlighted in grey. Within the ATPase domain the catalytic center, comprising Walker A, Walker B and motif C, are highlighted in green. SFX is located close to the Walker B domain and is colored blue. (B) Zoomed in view of the SFX binding pocket showing the interacting residues and electron density for the bound drug. Green residues are part of the Walker B motif. (C) Protein-Ligand Interaction profile of SFX with the 2C residues. Hydrophobic interactions are shown as dashed lines (P159, M175, L178, P182 and D186). Hydrogen bonds are shown as blue lines (C179) and water mediated hydrogen bonds are colored light blue (D186). Figure generated using PLIP^31^. (D) Overlay of the 2C crystal structure of CV-B3 (grey) with the 2C crystal structure of EV-A71 (PDB: 5GRB, chain A) (blue) in complex with ATP-γ-S.

The accessibility of SFX can be associated with two main conformational differences between the structure of EV-A71 and CV-B3 2C proteins. The first is a 10° tilt of the α2 helix (Figure 1D), enlarging the binding site of SFX by moving D186 away from the methylamine group of SFX. Together with this, the Walker B-carrying loop is positioned away from the binding pocket in the CV-B3 Δ116-2C, as illustrated by the 210° rotation of the side chain of Q180, making the cavity accessible to SFX. In addition, residues 180-182, located downstream of the Walker B motif, participate in main chain hydrogen bonding with the 224-AGSINA-229 loop, located downstream of motif C (Figure 2A). This provides direct evidence for crosstalk between the SFX binding site and the AGSINA loop which is a known hotspot for compound resistance and dependence mutations^18^.

**Figure 2.**
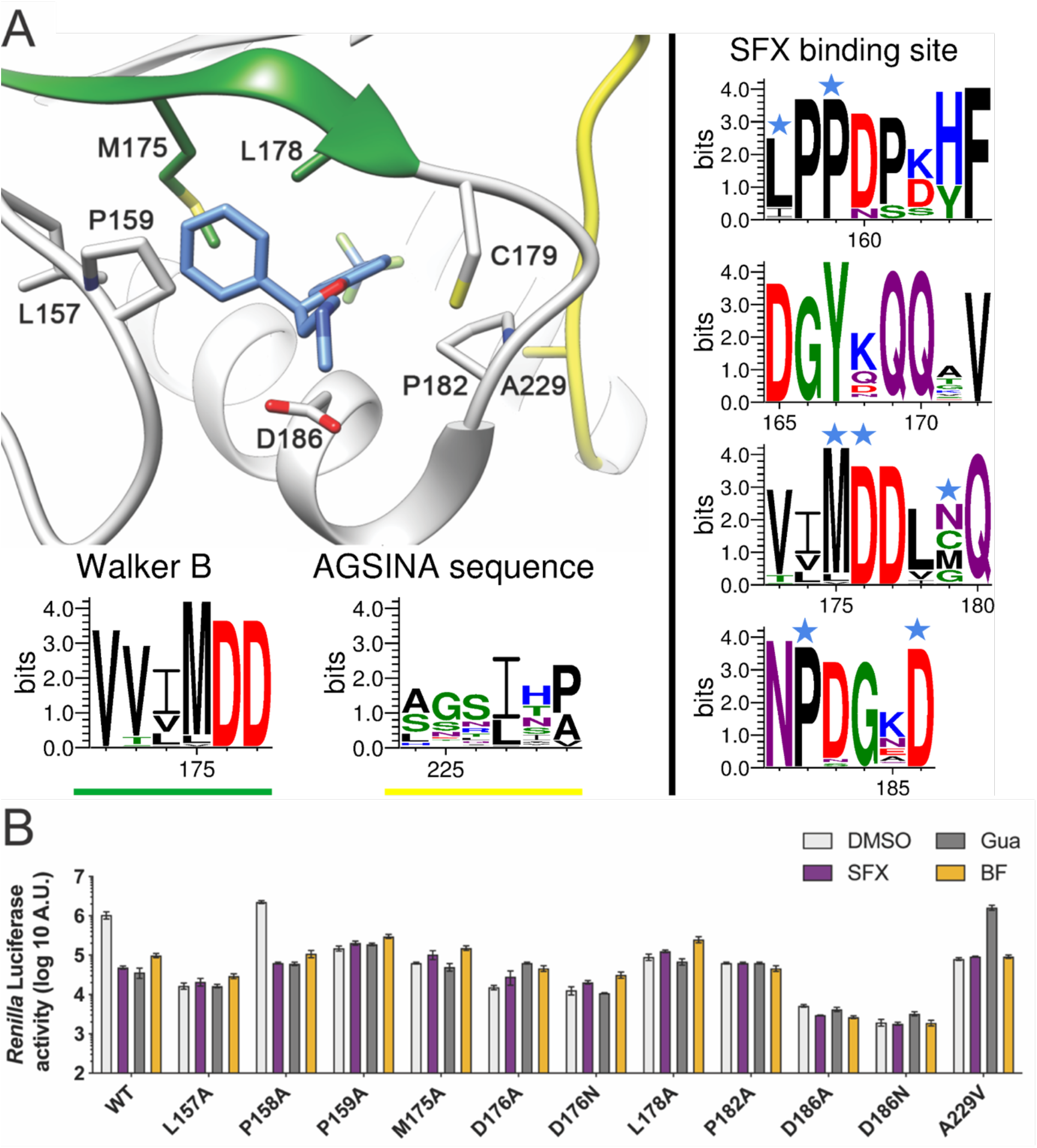
Mutational analysis of the SFX binding site in CV-B3 2C. (A) Schematic structure of CV-B3 2C highlighting the residues involved in SFX binding. The Weblogo represents the conservation of amino acids based on an alignment of the 2C protein from EVs which cause disease in humans. Virus sequences which were used for the alignment can be found in Supplement Part XYZ. The blue stars on top of the amino acid residues indicate an interaction with the ligand SFX. (B) The residues which interact with SFX were introduced into an infectious CV-B3 cDNA clone containing the *Renilla* luciferase reporter gene upstream of the capsid coding region (Rluc-CV-B3). In vitro transcribed RNA was transfected into cells and Renilla luciferase was used as a sensitive and quantitative read out for viral RNA replication.

### Mutational analysis of the SFX binding site

Sequence analysis of the SFX binding site revealed that all residues, except cysteine at position 179, are conserved in SFX resistant and sensitive EV species (Figures 2A and S3). To assess the functional consequences of alterations in the SFX binding residues, we performed mutagenesis of amino acids within, or adjacent to, the drug binding pocket. L157A, P158A, P159A, M175A, D176A, D176N, L178A, P182A, D186A and D186N were reverse engineered into a CV-B3 infectious cDNA clone containing a Renilla luciferase (CV-B3-Rluc) reporter gene. The replication efficiency of these mutants was then assessed in the presence or absence of 2C inhibitors. Nearly all substitutions abrogated viral RNA (vRNA) replication. Only P158A mutant, which is not in direct contact with SFX, was able to replicate. This mutation did not alter the sensitivity of 2C to SFX or GuaHCl (Figure 2B). We next tested the impact of introducing less stringent mutations into the SFX binding site. These mutations were selected using sequence information from other picornaviruses such as hepatitis A virus and Aichivirus (Figure S3). Surprisingly, even the introduction of chemically similar amino acids (i.e. L178I and D186E) resulted in no detectable vRNA replication (Figure S4). Consistently, upon introduction of mutations into a CV-B3 infectious clone without a reporter gene, no viable viruses were recovered, except for mutation P158A. Notably, after independent passages of cells transfected with the P159A mutant, viruses were recovered carrying a second-site compensatory mutation, namely A229V, which lies in proximity to Pro159. This substitution was previously shown to confer dependence to several compounds, including GuaHCl, HBB and MRL-1237^18,20,22,32^. Virus carrying the P159A/A229V double mutation showed a small decrease in sensitivity to SFX (Figure S5). Taken together, these data demonstrate that residues in the SFX binding pocket are highly mutationally constrained, presumably to preserve the function of 2C.

To gain more insight into the binding pocket of SFX, we raised resistant CV-B3 via a clonal selection procedure as described previously^29^. Several SFX-resistant viruses were obtained, containing single (I227V), double (I227V/A229V) or triple (A224V/I227V/A229V) mutations in the 224-AGSINA-229 loop (Figure S6). Mutations in this loop have been previously identified in CV-B3 resistant to 2C-targeting compounds^18,33^. In addition, we previously demonstrated that substitutions C179Y, C179F and F190L also provide resistance to SFX^29^. Notably, these latter mutations, as well as 224-AGSINA-229 loop mutations, provide cross-resistance against dibucaine (Fig S7). Our crystal structure of CV-B3 Δ116-2C in complex with SFX now allows us to cluster these mutations into two groups, those directly involved in SFX binding site (i.e. C179F, C179Y and F190L), and those in the 224-AGSINA-229 loop located downstream of the Walker C motif, which do not directly interact with SFX (Figure 2). The exact role of the 224-AGSINA-229 loop in conferring drug resistance, or dependence, is difficult to assess. However, its contribution to stabilization of the Walker B-containing loop in an “open conformation”, suitable for SFX binding, corroborates this motif as a credible hotspot for resistance mutations. To further demonstrate that this loop plays an important role in fluoxetine sensitivity, we transplanted the 224-AGSINA-229 loop from CV-B3 2C into the 2C protein of the EV-A71 BrCr strain, which contains 224-ASNIIV-229 and is insensitive to SFX, despite the SFX binding residues being conserved. Indeed, EV-A71 harboring the 224-AGSINA-229 loop gained sensitivity towards SFX, providing evidence that the 224-AGSINA-229 loop contributes to drug sensitivity (Figure S8).

### Engineering an enzymatically active 2C hexamer to study the mechanism of action of SFX and other inhibitors

The 2C protein contains three canonical motifs required for nucleotide triphosphate binding and hydrolysis, Walker A, Walker B and motif C^11^. Previous experiments demonstrated that SFX has little effect on the ATPase activity of the truncated, monomeric 2C protein of CV-B3^33^. However, the optimal activity of SF3 helicases requires their ATPase domains to be arranged in a hexameric complex^30,34,35^. Therefore, studies of truncated 2C proteins should be complemented with biochemical analysis of the enzymatically active hexameric complex. It has been demonstrated previously that the N-terminal domain of 2C is important for oligomerization and ATPase activity, but also renders the protein poorly soluble^13,14,19^. With this in mind, we attempted to purify the MBP-tagged full-length CV-B3 2C, which resulted in a heterogenous protein preparation as shown by the size exclusion chromatography gel filtration profile. This indicated that a mixture of high molecular weight 2C complexes were present, including a putative hexamer peak (Figure S9). As an alternative strategy, we sought to uncouple CV-B3 2C from the unfavorable biochemical properties of the N-terminal amphipathic helix but retain the ability to form hexamers. Fusion of N or C-terminal ‘assistant hexamer’ proteins is an established method to study AAA+ ATPases^36–38^. Based on the proposed hexamer model described for EV-A71 and PV 2C^16,17^, we hypothesized that we could fuse a hexameric parallel coiled-coil, CC-hex-D24^39^, to the N-terminus of CV-B3 Δ116-2C via a 10 amino acid linker sequence (Figure 3A-B). This would, in principle, enhance the local concentration of the 2C protein and correctly orientate the molecules in a manner reminiscent of membrane attachment, normally facilitated by the N-terminal amphipathic helix^40^. The small size of the CC-hex coiled-coil (32 amino acids) also permits the additional attachment of a cleavable His-MBP tag to promote solubility and facilitate affinity purification of the protein (Figure S10). As expected, the hexΔ116-2C elutes as a single peak which corresponds to a hexamer whereas Δ116-2C is exclusively monomeric (Figure 3C). We next employed the malachite green assay to assess the ATPase activity of our hexΔ116-2C, which demonstrated that the engineered protein is enzymatically active. In contrast, the monomeric protein and catalytic dead D177A mutant showed little to no ATPase activity (Figure 3D and S11). Altogether, the data show that the hexamerization is necessary to promote the 2C-dependent ATPase activity, offering a technological platform to screen or validate 2C ATPase inhibitors.

**Figure 3.**
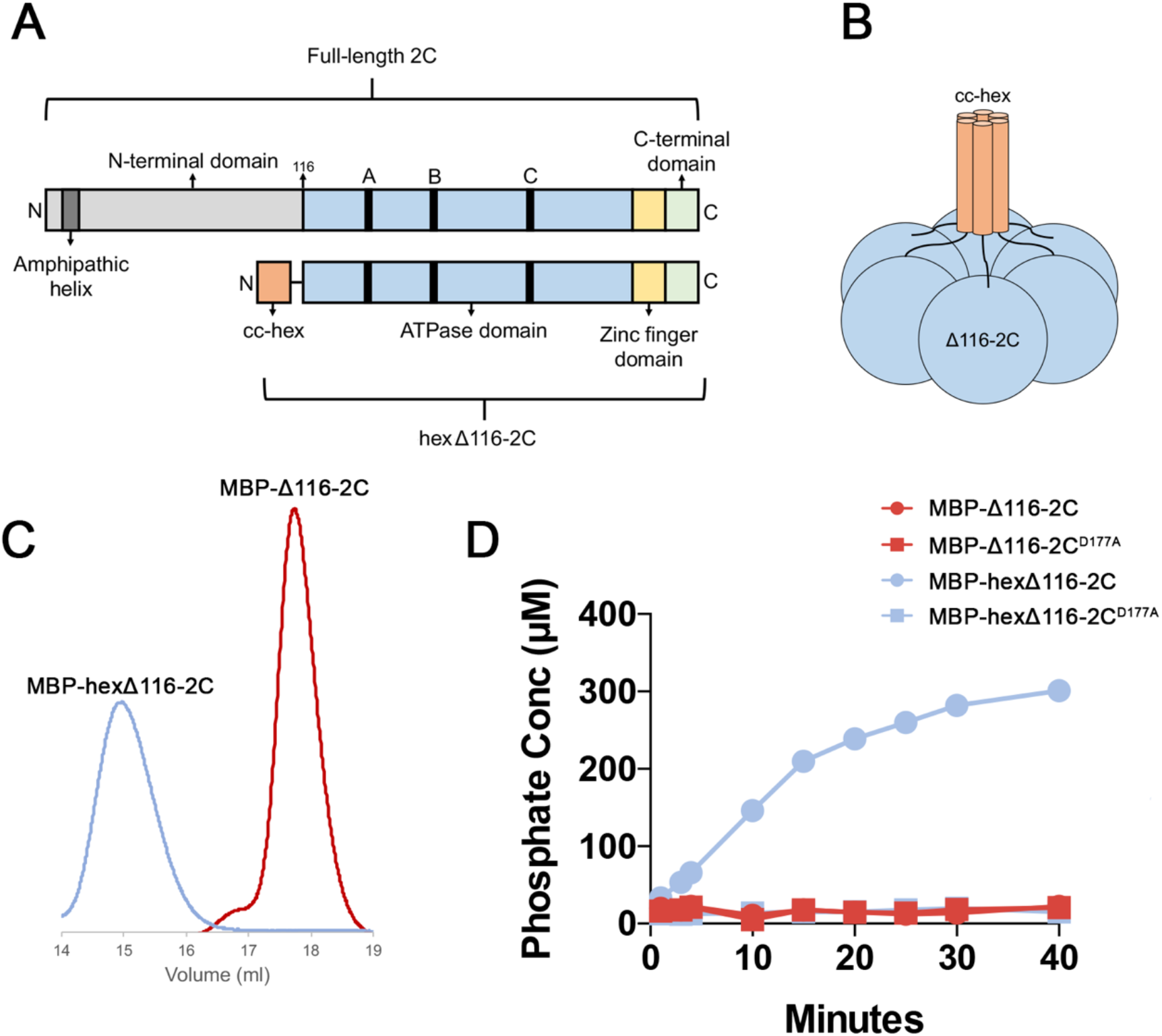
Engineering a soluble, enzymatically active 2C hexamer. (A) Linear representation of the full-length CV-B3 2C protein (top) and the engineered hexΔ116-2C construct (bottom). The full-length 2C protein comprises an N-terminal domain (light grey), ATPase domain (blue), Walker A, B and C motifs (black), zinc finger domain (yellow) and C-terminal extension (green). In hexΔ116-2C (residues 116-329), the N-terminal domain, which contains the amphipathic helix (dark grey), is replaced by the 32-residue cc-hex coiled coil sequence (orange). (B) Schematic representation of the hexΔ116-2C construct. (C) Migration profile of the MBP tagged monomeric Δ116-2C and hexΔ116-2C by size-exclusion chromatography. (D) Comparison of the ATPase activity for the WT hexΔ116-2C, the hexΔ116-2C D177A mutant, WT Δ116-2C and Δ116-2C D177A mutant.

Next, we investigated the effect of the fluoxetine enantiomers - SFX and RFX - on 2C-dependent ATP hydrolysis (Figure 4A). We showed that SFX inhibits ATPase activity in a dose dependent manner with an EC50 value of ∼5 µM, in the same concentration range as the dissociation equilibrium constant (Kd of 9.5 μM) obtained in a binding assay^26^. Consistent with the binding assay, RFX was markedly less efficient at inhibiting ATPase activity. We also tested the inhibitory effect of dibucaine, HBB and GuaHCl, known or suspected to target the 2C protein^19,21,24^. All of the compounds reduced the ATPase activity of 2C in a dose-dependent manner, underscoring the link between inhibition of ATP hydrolysis and antiviral effect (Figure 4B-C).

**Figure 4.**
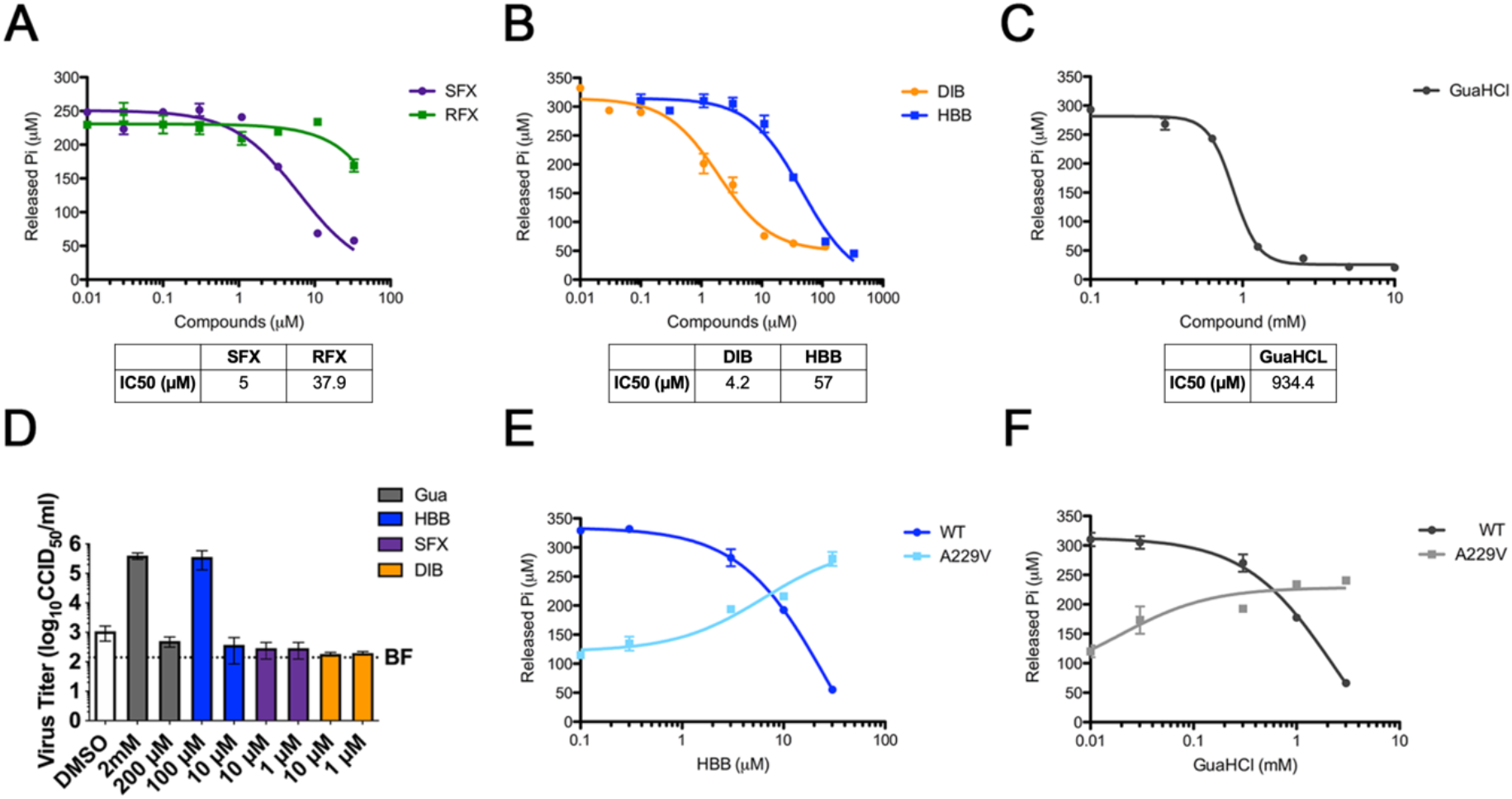
2C inhibitors reduce the ATPase activity of 2C and have antiviral effect. (A-C) Effects of SFX and RFX (A), dibucaine and [2-(alpha-hydroxybenzyl)-benzimidazole] (HBB) (B), or guanidium hydrochloride (GuaHCl) (C) on 2C-dependent ATPase activity. (D) Replication of CV-B3 containing the A229V 2C mutation in the presence of GuaHCL, HBB, SFX or DIB (or a non-related replication inhibitor, BF, which targets a host lipid kinase). At 8 hours post-infection, cells were freeze-thawed and virus titers of lysates were determined. (E, F) Comparison of ATPase activity of WT 2C and the A229V mutant in the presence of HBB (E) and GuaHCL (F).

**Figure 5.**
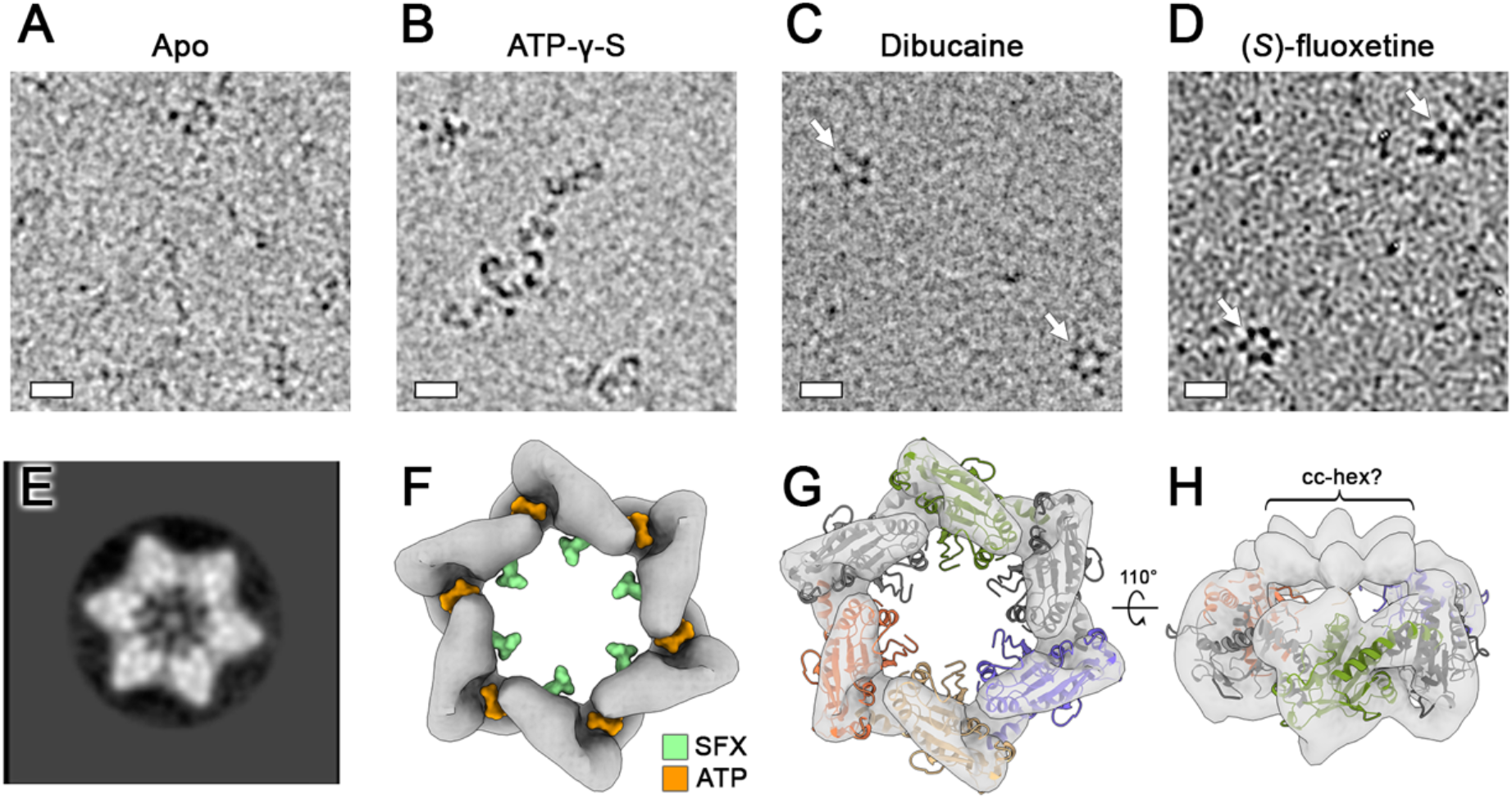
SFX and dibucaine stabilize the 2C hexamer. (A) Cryo-electron micrographs of apo, (B) ATP-y-S incubated, (C) dibucaine incubated or (D) SFX incubated hexΔ116-2C (scale bar = 10 nm). (E) Representative 2D class average from untilted micrographs of SFX incubated hexΔ116-2C. Weak density can be observed within the central of the 2C hexamer. (F) Surface representation of the 12 Å resolution cryo-EM map of hexΔ116-2C in complex with SFX, generated from 30° tilted cryo-EM data. The map is shown at a contour level of 0.0661 and the binding positions of SFX (green) and ATP (orange) are superposed. (G) As shown in (F) with a hexameric 2C model, generated using the crystal structure of the monomer, overlaid. For clarity, pore-loop residues 85-100 are omitted. (H) A 110° rotated view of the model shown in (G), with the map shown at a contour level of 0.059. Putative density for the linker and cc-hex is indicated.

It is known that besides drug-resistance, 2C targeting compounds can induce drug-dependence mutations. In the latter scenario, the presence of the compound is required for efficient enterovirus replication^18,20,32,35^. The 2C mutant, A229V, has been shown to confer dependence on GuaHCl for efficient CV-B3 replication^18^, but the underlying mechanism is unknown. Upon testing replication of CV-B3 carrying the A229V 2C mutation, we observed efficient replication in the presence of GuaHCl and HBB, but not SFX and dibucaine (Figure 4D). When the A229V mutant was introduced into our hexΔ116-2C construct, we observed reduced specific ATPase activity compared to the WT protein (Figure 4E). Remarkably, the level of ATPase activity observed with the WT 2C protein is fully and partly recovered in the presence of HBB and GuaHCl, respectively (Figure 4E-F). These data reveal that the A229V mutation negatively impacts the ATPase activity of 2C and that GuaHCl and HBB can restore the enzymatic activity.

### SFX and dibucaine stabilize the 2C hexamer

Having demonstrated that SFX and dibucaine can inhibit the 2C-dependent ATPase activity, we next wanted to determine how these compounds affect the quaternary structure of 2C. Cryo-electron microscopy analysis of our untagged engineered protein (Figure S12), in the absence of any nucleotide or inhibitor, did not reveal any distinct oligomeric structures (Figure 4A). When the non-hydrolysable ATP analogue, ATP-γ-S, was added, ‘hook-shaped’ oligomers were apparent (Figure 4B). When hexΔ116-2C was incubated with SFX or dibucaine, well-defined hexamers were discernable (Figure 4C-D). Single-particle analysis of SFX bound hexamers produced distinct star-shaped 2D class averages, some of which contained an electron dense feature in the central channel (Figure 4E). Efforts to determine the cryo-EM structure of the SFX-bound 2C hexamer were hampered by strong preferred orientation, low abundance, and instability of the particles. Nevertheless, a ∼12 Å resolution reconstruction of the complex was determined by employing a 30° stage tilt during data collection (Figure 4F and S13-14). The cryo-EM map corresponds to ∼140 kDa of ordered molecular weight and has a diameter of ∼110 Å, consistent with the dimensions of the EV-A71 2C hexamer model proposed previously^16^. Moreover, a cleft is present between the 2C protomers, which correlates with the expected location of the ATP binding pocket (Figure 4G). We next generated a hexameric model of the SFX-bound CV-B3 2C, based on the crystal structure of the JC polyomavirus large T antigen^41^, as described by Guan et al^16^, and fitted this as a rigid-body into our cryo-EM map. The overall agreement between the cryo-EM density and the hexameric model of CV-B3 2C supports the proposed mechanism of C-terminus mediated oligomerization. Indeed, in our crystal structures, the C-terminal helix (α6, α7) which appear to be broken in a “door knob” fashion into two helices α6 and α7, due to crystal packing, and α7 helix is in intensive contact with the Zinc binding domain of a symmetrical chain (Figure S15).

Taken together, these results indicate that SFX or dibucaine binding locks 2C in a hexameric state which does not permit ATP hydrolysis and, by extension, the associated functions such as helicase activity. Consistent with our cryo-EM observations, when size exclusion chromatography is performed on hexΔ116-2C in the presence of SFX, the complex exhibits an increased retention time, indicative of a reduced hydrodynamic radius (Figure S16A). In contrast, the presence of SFX did not greatly alter the elution profile of the monomeric Δ116-2C protein, except for a slightly more pronounced left shoulder peak, indicative of low-level oligomerization (Figure S16B). Therefore, it is likely that the SFX-mediated stabilization of the hexamer requires not only a high local concentration of protein, but also their correct orientation with respect to one another.

## Discussion

Enteroviruses are globally prevalent pathogens responsible for many diseases. There is an unmet need for broad-spectrum therapeutics to treat EV infections. The EV 2C protein is an attractive target for direct-acting antivirals, given that it is a highly conserved and functionally indispensable protein, undertaking several pleotropic functions during the viral lifecycle including membrane rearrangement, RNA unwinding and genome encapsidation^10^. Over the last decades, several structurally disparate compounds targeting 2C, such as the FDA-approved drugs fluoxetine and dibucaine, amongst others, were identified^23,24^. Subsequently, a number of resistance mutations were obtained to gain insights into their mode of action^33^. However, the binding sites and the mode of action remained elusive, hampering the rational-based improvement of the compounds. It has often been hypothesized that 2C inhibitors bind directly in the catalytic site. One study suggested that dibucaine analogues target the ATP binding site^42^, but no direct evidence supporting this hypothesis was presented. Recently, we obtained evidence for an allosteric binding site based on CV-B3, EV-A71 and EV-D68 mutants that were raised against a novel and highly potent 2C inhibitor^29^. Viruses resistant to this inhibitor contained mutations in, or adjacent to, the α2 helix, which is distal to the catalytic site. Here, we provide a high-resolution crystal structure of the CV-B3 2C protein in complex with SFX. This crystal structure revealed a hydrophobic binding pocket that accommodated the ligand between the α2 helix and the Walker B domain. Our data prove that SFX indeed targets an allosteric site on 2C that is clearly distinct from the catalytic site. The obtained structure not only provides valuable insights into the mode of action of SFX but also represents a novel tool for structure-based drug design of highly potent 2C inhibitors.

Several structural features are associated with the accessibility of SFX to bind the 2C protein of CV-B3 but not other enteroviruses. One structural key determinant allowing SFX to access the binding pocket is the 224-AGSINA-229 loop which is also a hot spot of resistance mutations. This is supported by our finding that exchange of the 224-ASNIIV-229 loop of the SFX insensitive EV-A71 BrCr with the 224-AGSINA-229 loop implemented a SFX sensitive phenotype to the resulting chimeric EV-A71 virus. Clearly, the 224-229 loop is not the only determinant for SFX sensitivity since we discovered previously that several clinical isolates of EV-A71 without changes in the 224-ASNIIV-229 are sensitive to SFX. Unfortunately, and despite extensive efforts, an apo-structure of CV-B3 is not available to elucidate the structural changes which occur upon SFX binding. Comparison of the SFX-bound CV-B3 2C to that of EV-A71 (apo and ATP-γ-S-bound) and PV (apo) shows that the α2-helix is tilted 10° away from the SFX binding site in CV-B3 2C, which creates the solvent accessible volume required for drug binding. Similarly, the Walker B loop of CV-B3 2C is positioned further away from the SFX binding site compared to EV-A71 and PV 2C, which further contributes to formation of the hydrophobic pocket required to accommodate SFX. This suggests that, in solution, 2C exhibits a high degree of conformational plasticity and that SFX bind to and stabilizes, an “open conformation” of 2C. Ultimately, an apo structure of CV-B3 2C is required to confirm this hypothesis.

The crystal structure of the 2C monomer in complex with SFX is not sufficient to decipher its mode of action as this should take into account the expected hexameric nature of AAA+ proteins. Several lines of evidence suggest that the catalytically active form of 2C is composed of higher oligomeric structures^12,13,35^. Production of 2C without its N-terminal helical domain yields monomeric protein that has little, if any, ATPase activity^33^. To overcome the biochemical limitations of 2C and to study the mode of action of 2C-targeting antivirals, we engineered an ATP hydrolyzing, hexameric form of the 2C ATPase domain. The resulting construct produced homogenous oligomers that displayed robust ATPase activity compared to the inactive monomeric form of 2C (Figure 3). Using the engineered protein, we demonstrate that the FDA approved drugs fluoxetine, dibucaine as well as GuaHCl and HBB inhibit the ATPase activity in a dose-dependent manner. Thus, our hexameric 2C construct represents a novel tool for *in vitro* screening of 2C-targeting ATPase inhibitors. This tool might be of particular interest to identify potential 2C inhibitors for difficult-to-culture enteroviruses such as the RV-C species members^43^. Besides this, this platform can provide mechanistic insights into the mode of action of specific mutations, as shown here for the drug-dependent A229V mutation which we revealed to impair ATPase activity in the absence of inhibitors, but to stimulate ATPase activity in the presence of GuaHCl and HBB.

A hexameric structure of 2C has been long sought-after in the picornavirus field but has remained elusive due to the unfavorable biochemical properties of the full-length protein, especially its amphipathic N-terminal α-helix. Therefore, cryo-EM analysis of CV-B3 2C protein, incubated with SFX, provides the first 3D reconstruction of a hexameric complex, and represents a significant step towards understanding this enigmatic protein. Although artificially generated by the addition of an hexamerization domain, the recovery of the ATPase activity and the binding of SFX unambiguously confirm the relevance of the construct. Previous studies have shown that the 2C protein of foot-and-mouth disease virus (FMDV), a member of the *Aphthovirus* genus within the picornavirdae family, employs a coordinated ATP hydrolysis mechanism. Using negative stain analysis, the same study demonstrated that FMDV 2C(34-318), containing the motif C mutation N207A (which interferes with ATP hydrolysis but not binding), could form hexamers in the presence of ATP and RNA^14^. It is likely that enterovirus 2C proteins must undergo conformations changes during ATP hydrolysis, RNA binding and unwinding, to drive their helicase function. Our biochemical and cryo-EM analysis suggests that SFX and dibucaine binding prevents propagation of these conformational changes by locking 2C in a defined hexameric state. A similar mechanism was shown for a small molecule inhibitor of the human AAA+ ATPase, p97^44^. The allosteric p97 inhibitor binds at the interface of two adjacent AAA domains and prevents the conformational changes that are required for ATPase activity of p97. As SFX does not bind to a 2C-2C interface, it’s likely that drug binding induces conformation changes within the monomer that translate to stabilization of the hexamer. As shown for several AAA+ ATPases, cryo-EM provides insights that are not available from crystallographic analysis alone^44–46^. In the case of 2C, the observed stabilizing effect of SFX, observed in solution, complements the high-resolution information obtained by crystallography and provides insights into the allosteric inhibitory mechanism of 2C targeting compounds. Our findings emphasize the importance of using an integrative structural biology approach to fully understand the function and mechanisms of inhibition of 2C. Previous studies have shown that purified, full-length poliovirus 2C cannot be uncoupled from its solubility tag, and contains a mixture of oligomeric species – both of which would hamper structural analysis^13^. Similarly, an N-terminally extended 2C protein of echovirus 30 was shown to form heterogenous ring-shaped structures^15^. Here we demonstrate that replacement of the N-terminal region of 2C with a hexamerization domain can produce a homogenous complex amenable to cryo-EM. Going forward, our engineering strategy should facilitate further structural and functional studies of oligomeric 2C proteins, and their interaction with inhibitors, viral RNA, and other viral/host factors.

## Material and methods

### Cells and reagents

Buffalo Green Monkey (BGM) cells (purchased from European Cell Culture Collection [ECACC]) and HeLa R19 cells (American Type Culture Collection [ATCC]) were cultured in Dulbecco’s Modified Eagle Medium (DMEM; Lonza, Switzerland) supplemented with 10% (vol/vol) fetal calf serum (FCS; Lonza). All cell lines were grown at 37 °C in 5% CO2. Medium was refreshed every 2–4 days, and cells were passaged at >80% confluence with the use of PBS and trypsin-EDTA (0.05%) for up to 10 passages. The cell lines were routinely tested for mycoplasma contamination. SFX, RFX, GuaHCl and dibucaine were purchased from Sigma Aldrich. HBB was provided by Prof. Andrea Brancale (Cardiff University). GuaHCl and ATP-γ-S ((Sigma Aldrich) were dissolved in water at 2 M and 20 mM stock concentration, respectively. All other compounds were dissolved in DMSO at 10 mM stock concentration.

### Cloning and construct design

For crystallography experiments, the coding sequence of CV-B3 2C (amino acids 117 to 329) was cloned into the expression vector pmCox 20A as previously described^47^. For ATPase assays and cryo-EM experiments, the coding sequence of CV-B3 2C (amino acids 116 to 329) was inserted into pET-28b plasmid. The hexΔ116-2C construct was produced by inserting a 32-residue codon optimized cc-hex coding sequence followed by a linker (resulting peptide: GELKAIAQELKAIAKELKAIAWELKAIAQGAG; linker: GSGSYFQSNA Genscript) at the 5’ of the CV-B3 2C coding sequence. For both constructs, a 3C protease cleavable N-terminal His6-MBP tag was included to facilitate expression and purification. An overview of the hexΔ116-2C construct is shown in Figure S10. The D177A and A229V variants were produced by site directed mutagenesis using the Δ116-2C and hexΔ116-2C plasmids as a template.

### Protein expression and purification

The CV-B3 Δ116-2C protein used for crystallogenesis was produced in *Escherichia coli* T7 Iq Express pLysS (New England BioLabs). Cells were grown in TB medium containing 100 µg/mL of ampicillin and 34 µg/mL chloramphenicol at 37°C. Expression was induced overnight at 17°C with 0.5 mM isopropyl -thiogalactopyranoside (IPTG). The cells were harvested by centrifugation and resuspended in lysis buffer (50 mM Tris, 300 mM NaCl, 10 mM imidazole, 5% glycerol, 0.1% Triton) complemented with 0.25 mg/mL lysozyme, 10 µg/mL DNase and 1 mM of PMSF. The bacterial sample was sonicated and centrifuged at 18000 g for 30 min 4°C. Soluble CV-B3 Δ116-2C was purified from the supernatant by immobilized metal-affinity chromatography (IMAC) on a HisTrap column (GE Healthcare) and eluted with a buffer consisting of 50 mM Tris, 300 mM NaCl, 500 mM imidazole pH 8. The eluted protein was subject to a dialysis in 50 mM Tris pH 8, 400 mM NaCl,10 mM Imidazole, 1 mM DTT (dithiothreitol), followed by an overnight cleavage by TEV protease (molar ratio: 1/20), followed by a second IMAC to remove the tag and uncleaved protein. The recombinant protein was finally purified by size-exclusion chromatography (SEC) on a HiLoad 16/60 Superdex 75 column (GE Healthcare) in 10 mM Hepes pH 7.5 and 300 mM NaCl.

For electron microscopy and functional assays, *Escherichia coli* Rosetta 2 (DE3) Cells (Sigma-Aldrich) were transformed with the plasmids coding for the WT, D177A and A229V CV-B3 2C proteins and grown in 2xYT medium containing 50 µg/mL of kanamycin and 30 µg/mL chloramphenicol at 37°C until the OD600nm reach 0.3. The temperature was then reduced to 18°C and when the OD600nm reached 0.5, protein expression was induced with 0.5 mM isopropyl -thiogalactopyranoside (IPTG). Following expression for 16 hours, the cells were harvested by centrifugation and resuspended in lysis buffer (50 mM Tris, 300 mM NaCl, 10 mM imidazole, 5% glycerol, 0.1% Triton, 1 mM MgCl2) complemented with 0.25 mg/mL lysozyme, 10 µg/mL DNase I and 1 EDTA-free protease inhibitor cocktail. The bacterial sample was sonicated and centrifuged at 25000 g for 45 min at 4°C. The supernatant was passed through a 0.45 µM filter and incubated with 1 mL of Ni-NTA resin at 4°C for 1 hour on a roller. The beads were washed with wash I buffer (50 mM Tris, 300 mM NaCl, 10 mM imidazole, 1 mM MgCl2) and wash II buffer (50 mM Tris, 300 mM NaCl, 10 mM imidazole, 1 mM MgCl2) and then eluted with a buffer consisting of 50 mM Tris, 300 mM NaCl, 500 mM imidazole pH 8. For ATPase assays, the eluted proteins were concentrated to ∼200 µL and purified by size-exclusion chromatography on a Superose 6 10/300 GL column (GE Healthcare) in 10 mM Hepes pH 7.5, 150 mM NaCl and 10 mM MgCl2. The protein-containing fractions were pooled, concentrated to ∼4 mg/mL and flash-frozen in 20 µL aliquots. To remove the N-terminal His6-MBP tag for cryo-EM analysis, the Ni-NTA eluted hexΔ116-2C protein was concentrated to 200 µL, combined with 2 µg of HRV-3C protease (Sigma) and dialyzed overnight in buffer containing 50 mM Tris (pH 8), 200 mM NaCl and 0.5 mM DTT. The cleaved protein was finally purified by size-exclusion chromatography (SEC) on a Superose 6 10/300 GL column (GE Healthcare) in 25 mM Tris (pH 8), 300 mM NaCl, 1 mM MgCl2. The fractions corresponding to hexΔ116-2C were pooled, concentrated to ∼4 mg/mL and flash-frozen in 10 µL aliquots.

### Crystallogenesis, data collection and structure determination

Structure Screens 1 and 2 and Stura Footprint Screen (Molecular Dimensions Ltd) were used for initial screening. Trials were assessed in SWISSCI 3 Lens Crystallization Plates with three wells per reservoir using 400, 300 and 200 nL drops. The drops contained increasing volumes (100, 200 and 300 nL) of protein solution at a concentration of 8-9 mg/mL in complex with or without 2.5 mM SFX (from 50 mM stock in 100% DMSO) and 100 nL mother liquor. Crystal hits were observed only in presence of *S*FX, in 0.1M MES pH 6.5, Polyethylene glycol (PEG) 20K 12%. Optimization of crystal-growth conditions led to the following condition: 0.1M MES pH 5.7 – 6.7, PEG 20K 7.5/15 %. Crystals appeared after 24 h. They were soaked in 25% (*v*/*v*) PEG 200 added to the mother liquor and cooled in liquid nitrogen. X-ray diffraction data were collected at European Synchrotron Radiation Facility, Grenoble (ESRF). The structure was solved by molecular replacement using Phaser with PV-2C-ΔN-3Mut structure (PDB code: 5Z3Q, D chain) as the searching model and was refined to a resolution of 1.4Å. The crystal belongs to a space group of P212121, with 1 copy of CV-B3 Δ116-2C in the asymmetric unit (ASU). In addition, crystals were also obtained in another crystallization solution containing 0.1M MES pH 6.3 – 6.7, PEG 5 000 MME 23/33 %, 0,2M ammonium sulphate. Crystals in mother liquor were soaked with 0.3 µL of 20 mM ATP overnight at 20°C. The datasets were collected at Proxima-2A-Synchrotron SOLEIL. The structure was solved by molecular replacement using Phaser with the structure of CV-B3 Δ116-2C in complex with SFX as a search model and refined to a resolution of 1.5 Å. All of the data sets were processed using XDS^48^ and scaled with *SCALA* (Collaborative Computational Project, Number 4, 1994). The model building was performed in Coot^49^, and the structures were refined using phenix.refine^50^. The statistics of data collection, structure refinement and structure validation were summarized in Table S1. Figures of CV-B3 Δ116-2C structures and structural alignments were drawn by Chimera^51^.

### Protein stability assay

To assess the folding of CV-B3 Δ116-2C and its ability to bind SFX, the thermal stability of recombinant CV-B3 Δ116-2C in presence of SFX was monitored by fluorescence-based thermal shift assay (TSA) using a Bio-Rad CFX Connect. TSA plates were prepared by dispensing into each well the 2C protein (final concentration of 15 μM in10 mM Hepes pH 7.5 and 300 mM NaCl) which was mixed with 1 μL of SFX (from 20 mM stock in 100% DMSO, 1 mM final concentration in 4% DMSO) and a SYPRO orange solution in concentrations recommended by the manufacturer in a final volume of 25 μL.

### Reverse engineering of SFX interaction residues

The CV-B3 mutations in 2C L157A, P158A, P159A, M175A, M175I, M175G, D176A, D176N, L178A, L178I, P182A, D186A, D186E, D186N were first introduced into the Rluc-CV-B3 reporter viruses with Q5 site-directed mutagenesis kit (New England Biolabs, Bioké, Leiden, The Netherlands)^52,53^. The Rluc-CV-B3 reporter viruses contain a *Renilla* luciferase gene upstream of the capsid coding region which allows a fast and quantitative read-out for virus replication. Next, a 705 bp fragment containing the desired mutation was isolated using the enzymes BssHII and XbaI and reintroduced into the original non-mutagenized Rluc-CV-B3 backbone. After site-directed mutagenesis the newly generated plasmids were subjected to Sanger sequencing to confirm the existence of the 2C mutation. The corresponding primers which were used for side-directed mutagenesis can be found in Table S3. Viral RNA was transcribed *in vitro* using the T7 RiboMAX Express Large Scale RNA production system (Promega, Leiden, The Netherlands) according to the manufacturer’s protocol. In a 96-well, HeLa R19 cells were transfected with 7,5 ng of viral RNA. One hour after transfection, the medium was replaced by fresh medium and/or compound containing medium. After 8 hours, cells were lysed and luciferase activity was determined using the Renilla luciferase Assay System (Promega, Leiden, The Netherlands). Since most of the mutations abrogated virus replication, we cloned the above-mentioned 2C mutations into the p53CB3/T7 infectious clone using BssHII and XbaI^53^. Similar to the Rluc-CV-B3 plasmids, the infectious clones were linearized with MluI and RNA was transcribed *in vitro*, and 150 ng of viral RNA was transfected into 6-wells containing either HeLa R19 or BGM cells. All transfections were done in triplicates. Transfection of CV-B3-2C[A229V] was done in the presence of 1mM GuaHCl. 3 days after transfection, the transfected cells were subjected to three freeze-thaw cycles and the lysates were passaged 3 times on either HelaR19 or BGM cells, respectively. If CPE was observed during passaging of the viruses, viral RNA was isolated with the NucleoSpin RNA Virus kit (Macherey-Nagel, Leiden, The Netherlands) according to the manufacturer’s protocol. The viral RNA was reverse transcribed into cDNA with random hexamer primers using the TaqMan Reverse Transcription Reagents (Applied Biosystems) and PCR was performed to isolate the 2C regions with the forwards primer binding in 2B 5’ CTAACCAAATATGTGAGC 3’ and the reverse primer binding in 3A 5’ CTCACTGTCTACCGATTTGAG 3’. The same primers were used for Sanger sequencing of the PCR product. If virus was obtained, virus titers were determined by endpoint dilution titration and calculated according to the method of Reed and Muench and expressed as 50% cell culture infective dose (CCID50)^54^. The obtained CV-B3-2C[A229V] virus was titrated in the presence of 2mM GuaHCl.

### Single-cycle infection

Confluent HeLa R19 (25.000 cells/ well in a 96-well) cells were infected with virus at a multiplicity of infection (MOI) of 0.1 or 1 at 37 °C for 30 min. Next, the medium was removed, and fresh (compound-containing) medium was added to the cells. At the indicated time points, the medium was discarded, and cells were subjected to three times freeze thawing to determine the virus titers by endpoint dilution using the methods of Reed and Muench. In the case of RLuc-CV-B3 infection, cells were lysed 8 hrs post infection to determine the luciferase activity with the *Renilla* luciferase Assay System (Promega). The CV-B3-2C[A229V] mutant was titrated in the presence of 2mM GuaHCl.

### ATPase Assay

The release of inorganic phosphate during 2C-mediated ATPase hydrolysis was measured using the Malachite Green Phosphate Assay Kit (Sigma Aldrich, Zwijndrecht, The Netherlands). Each 50 µL reaction comprised 500 nM recombinant 2C protein and 1 mM ATP in reaction buffer (10 mM HEPES (pH 7.5), 150 mM NaCl and 10 mM MgCl2). The reaction buffer was always prepared fresh and recombinant 2C proteins were thawed immediately prior to use. For testing the inhibitory effect of 2C targeting compounds, the final DMSO concentration in each reaction was 1%. Samples were incubated for 30 min at 37°C and then diluted 4-fold with water to give a final ATP concentration of 0.25 mM, as per the manufacturer’s instructions. To terminate the enzyme reaction, 80 µL of the diluted sample was mixed with 20 µL of Solution AB in a 96-well plate. The plates were incubated for 30 min at room temperature and absorbance was measured using a microplate reader at OD630. To determine the phosphate concentration in each sample, the OD630 values were plotted against a standard curve using GraphPad Prism version 8.

### Cryo-electron microscopy

HexΔ116-2C (1 mg/mL) was either used directly for grid preparation or first incubated with 5 mM ATP-y-S, 100 µM SFX or 100 µM dibucaine (final DMSO concentration of 0.5%), at 4°C for 30 minutes. 3 μl of each sample was dispensed on Quantifoil R1.2/1.3 200-mesh grids (Quantifoil Micro Tools GmbH) that had been freshly glow discharged for 30 seconds at 20 mA using a PELCO easyGLow™ Glow Discharge Cleaning System (Tedpella). Grids were blotted for five seconds using Whatman No. 1 filter paper and immediately plunge-frozen into liquid ethane cooled by liquid nitrogen using a Vitrobot Mark IV plunger (Thermo Fisher Scientific) equilibrated to ∼95% relative humidity, 4°C. Screening of the apo, ATP-y-S incubated and SFX or dibucaine incubated samples was performed using a 200 kV Talos Arctica (Thermo Fisher Scientific) equipped with a Gatan K2 Summit direct detector. Subsequently, two data sets were collected on the SFX incubated sample using a Titan Krios Cryo-TEM (Thermo Fisher Scientific) operating at 300 keV. For the first data set, movies were collected using a K2 direct electron detector operating in electron counting mode, at 165,000× magnification corresponding to a calibrated pixel size of 0.842 Å/pix over a defocus range of −1.5 to −2.5 μm. 1527 movies were collected using a dose rate of 8.8 e^-^/pix/sec for a total of 4 seconds (40 fractions), resulting in a total exposure of ∼50 e^-^/Å^2^ (1.25 e^-^/Å^2^/fraction). To account for the preferred orientation exhibited by the 2C hexamers, an alpha tilt of +30 degrees was used for the second data collection. Movies were collected using a K3 direct electron detector operating in super resolution mode, at 64,000× magnification corresponding to a super resolution pixel size of 0.69 Å/pix over a defocus range of −2 to −4 μm. 6119 movies were collected using a dose rate of 12.6 e^-^/pix/sec for a total of 2 seconds (37 fractions), resulting in a total exposure of ∼54 e^-^/Å^2^ (1.45 e^-^/Å^2^/fraction). All cryo-EM data were acquired using the EPU 2 software (Thermo Fisher Scientific).

### Image processing

For the untilted data, collected movie stacks were manually inspected and then imported in Relion version 3.0.1^55^. Drift and gain correction were performed with MotionCor2^56^, and GCTF was used to estimate the contrast transfer function for each micrograph^57^. Approximately 200 particles were picked manually and 2D classified. The best resulting class was then used as a template for autopicking in Relion, resulting in 42,206 particles. Fourier binned (2 × 2) particles were extracted in a 240-pixel box and subjected to a round of 2D classification after which 3616 particles were retained. Using the ‘molmap’ command in UCSF chimera, residues 358-628 of the JC polyomavirus helicase (PDB ID: 5J40^41^) were used to generate a 50 Å resolution starting model. Particles selected from 2D classification were 3D auto-refined (with C6 symmetry), which produced a highly anisotropic map. Following unsuccessful attempts to solve the preferred particle orientation with buffer additives, tilted data collection was employed as described previously^58^. Tilted movies were Fourier binned (2 × 2) during motion correction with MotionCor2^56^, and goCTF was used to estimate the defocus gradient of each micrograph^59^. Approximately 1000 particles were picked manually and 2D classified. The best resulting class was then used as a template for autopicking in Relion, resulting in 180,069 particles. Fourier binned (2 × 2) particles were extracted in an 80-pixel box and subjected 3D classification, after which 53,854 particles were selected. After a second round of 3D classification, 11,024 particles were retained. A final round of no-alignment 3D classification was performed, yielding a final stack of 6856 particles. Subsequent 3D auto-refinement (with C6 symmetry) and post-processing yielded a map with an estimated resolution of 12Å, based on the gold-standard FSC=0.143 criterion. An ad-hoc negative B-factor of −200Å^2^ was applied during the final post-processing step. An overview of the data processing pipeline is shown in Supplementary Figure 13. Figures of the CV-B3 hexΔ116-2C reconstruction were made in UCSF ChimeraX^60^.

### Data availability

Atomic coordinates and structure factors for CV-B3 Δ116-2C in complex with SFX are deposited in the Protein Data Bank under accession codes 6S3A and 6T3W. The EM density map for the SFX incubated hexΔ116-2C has been deposited to the Electron Microscopy Data Bank under the accession EMD-12798. All reagents and relevant data are available from the authors upon request.

## Acknowledgments

DLH is funded from the European Union’s Horizon 2020 research and innovation program under the Marie Skłodowska-Curie grant agreement (No 842333) and holds an EMBO non-stipendiary long-term Fellowship (ALTF 1172-2018). PEK is the recipient of a scholarship of the Foundation “Méditerranée Infection”, Marseille. This work was supported by the European Union (Horizon 2020 Marie Skłodowska-Curie ETN ‘ANTIVIRALS’, grant agreement number 642434 to AB, FJMvK and BCo). This work is also supported by Netherlands Organization for Scientific Research (NWO-ECHO-711.017.002 to FJMvK) and NWO-VICI-91812628 to FJMvK), and from the Life Science Research Network Wales (grant no. NRNPGSep14008 to AB, an initiative funded through the Welsh Government’s Ser Cymru program). This work was supported by the European Research Council under the European Union’s Horizon2020 Programme (ERC Consolidator Grant Agreement 724425 - BENDER). This work benefited from access to the Netherlands Centre for Electron Nanoscopy (NeCEN) at Leiden University, an Instruct-ERIC center with assistance from Dr. Rebecca Dillard. We thank Dr. Mihajlo Vanevic for computational support.

## Notes

https://www.rcsb.org/structure/6S3A

https://www.rcsb.org/structure/6T3W

### Competing Interest Statement

The authors have declared no competing interest.

## References

1. Tapparel, C., Siegrist, F., Petty, T. J. & Kaiser, L. Picornavirus and enterovirus diversity with associated human diseases. Infection, Genetics and Evolution vol. 14 (2013).

2. Thibaut, H. J. et al. Toward antiviral therapy/prophylaxis for rhinovirus-induced exacerbations of chronic obstructive pulmonary disease: Challenges, opportunities, and strategies. Rev. Med. Virol. 26, (2016).

3. Chapman, N. M. & Kim, K. S. Persistent coxsackievirus infection: Enterovirus persistence in chronic myocarditis and dilated cardiomyopathy. Current Topics in Microbiology and Immunology vol. 323 (2008).

4. Puenpa, J., Wanlapakorn, N., Vongpunsawad, S. & Poovorawan, Y. The History of Enterovirus A71 Outbreaks and Molecular Epidemiology in the Asia-Pacific Region. Journal of Biomedical Science vol. 26 (2019).

5. Holm-Hansen, C. C., Midgley, S. E. & Fischer, T. K. Global emergence of enterovirus D68: A systematic review. The Lancet Infectious Diseases vol. 16 (2016).

6. Messacar, K. et al. Enterovirus D68 and acute flaccid myelitis—evaluating the evidence for causality. The Lancet Infectious Diseases vol. 18 (2018).

7. Aw-Yong, K. L., NikNadia, N. M. N., Tan, C. W., Sam, I. C. & Chan, Y. F. Immune responses against enterovirus A71 infection: Implications for vaccine success. Reviews in Medical Virology vol. 29 (2019).

8. Lin, J. Y., Kung, Y. A. & Shih, S. R. Antivirals and vaccines for Enterovirus A71. Journal of Biomedical Science vol. 26 (2019).

9. Baggen, J., Thibaut, H. J., Strating, J.R.P.M. & Van Kuppeveld, F. J. M. The life cycle of non-polio enteroviruses and how to target it. Nature Reviews Microbiology vol. 16 (2018).

10. Wang, S. H., Wang, K., Zhao, K., Hua, S. C. & Du, J. The Structure, Function, and Mechanisms of Action of Enterovirus Non-structural Protein 2C. Frontiers in Microbiology vol. 11 (2020).

11. Gorbalenya, A. E., Koonin, E. V. & Wolf, Y. I. A new superfamily of putative NTP-binding domains encoded by genomes of small DNA and RNA viruses. FEBS Lett. 262, (1990).

12. Xia, H. et al. Human Enterovirus Nonstructural Protein 2CATPase Functions as Both an RNA Helicase and ATP-Independent RNA Chaperone. PLoS Pathog. 11, (2015).

13. Adams, P., Kandiah, E., Effantin, G., Steven, A. C. & Ehrenfeld, E. Poliovirus 2C protein forms homo-oligomeric structures required for ATPase activity. J. Biol. Chem. 284, (2009).

14. Sweeney, T. R. et al. Foot-and-mouth disease virus 2C is a hexameric AAA+ protein with a coordinated ATP hydrolysis mechanism. J. Biol. Chem. 285, (2010).

15. Papageorgiou, N. et al. The 2C putative helicase of echovirus 30 adopts a hexameric ring-shaped structure. Acta Crystallogr. Sect. D Biol. Crystallogr. 66, (2010).

16. Guan, H. et al. Crystal structure of 2C helicase from enterovirus 71. Sci. Adv. 3, (2017).

17. Guan, H., Tian, J., Zhang, C., Qin, B. & Cui, S. Crystal structure of a soluble fragment of poliovirus 2CATPase. PLoS Pathog. 14, (2018).

18. De Palma, A. M. et al. The Thiazolobenzimidazole TBZE-029 Inhibits Enterovirus Replication by Targeting a Short Region Immediately Downstream from Motif C in the Nonstructural Protein 2C. J. Virol. 82, (2008).

19. Pfister, T. & Wimmer, E. Characterization of the nucleoside triphosphatase activity of poliovirus protein 2C reveals a mechanism by which guanidine inhibits poliovirus replication. J. Biol. Chem. 274, (1999).

20. Hadaschik, D., Klein, M., Zimmermann, H., Eggers, H. J. & Nelsen-Salz, B. Dependence of Echovirus 9 on the Enterovirus RNA Replication Inhibitor 2-(α-Hydroxybenzyl)-Benzimidazole Maps to Nonstructural Protein 2C. J. Virol. 73, (1999).

21. Klein, M., Hadaschik, D., Zimmermann, H., Eggers, H. J. & Nelsen-Salz, B. Picornavirus replication inhibitors HBB and guanidine in the echovirus-9 system: The significance of viral protein 2C. J. Gen. Virol. 81, (2000).

22. Shimizu, H. et al. Mutations in the 2C Region of Poliovirus Responsible for Altered Sensitivity to Benzimidazole Derivatives. J. Virol. 74, (2000).

23. Zuo, J. et al. Fluoxetine is a potent inhibitor of coxsackievirus replication. Antimicrob. Agents Chemother. 56, (2012).

24. Ulferts, R. et al. Screening of a library of FDA-approved drugs identifies several enterovirus replication inhibitors that target viral protein 2C. Antimicrob. Agents Chemother. 60, (2016).

25. Stark, P. & Hardison, C. D. A review of multicenter controlled studies of fluoxetine vs. imipramine and placebo in outpatients with major depressive disorder. J. Clin. Psychiatry 46, (1985).

26. Bauer, L. et al. Fluoxetine Inhibits Enterovirus Replication by Targeting the Viral 2C Protein in a Stereospecific Manner. ACS Infect. Dis. 5, (2019).

27. Manganaro, R. et al. Synthesis and antiviral effect of novel fluoxetine analogues as enterovirus 2C inhibitors. Antiviral Res. 178, (2020).

28. Musharrafieh, R. et al. Discovery of Quinoline Analogues as Potent Antivirals against Enterovirus D68 (EV-D68). J. Med. Chem. 62, (2019).

29. Bauer, L. et al. Rational design of highly potent broadspectrum enterovirus inhibitors targeting the nonstructural protein 2C. PLoS Biol. 18, (2020).

30. Hickman, A. B. & Dyda, F. Binding and unwinding: SF3 viral helicases. Current Opinion in Structural Biology vol. 15 (2005).

31. Salentin, S., Schreiber, S., Haupt, V. J., Adasme, M. F. & Schroeder, M. PLIP: Fully automated protein-ligand interaction profiler. Nucleic Acids Res. 43, (2015).

32. Eggers, H. J. & Tamm, I. Drug dependence of enteroviruses: Variants of Coxsackie A9 and ECHO 13 viruses that require 2-(α-hydroxybenzyl)-benzimidazole for growth. Virology 20, (1963).

33. Ulferts, R. et al. Selective serotonin reuptake inhibitor fluoxetine inhibits replication of human enteroviruses B and D by targeting viral protein 2C. Antimicrob. Agents Chemother. 57, (2013).

34. Gai, D., Zhao, R., Li, D., Finkielstein, C. V. & Chen, X. S. Mechanisms of conformational change for a replicative hexameric helicase of SV40 large tumor antigen. Cell 119, (2004).

35. Tolskaya, E. A. et al. Genetic studies on the poliovirus 2C protein, an NTPase A plausible mechanism of guanidine effect on the 2C function and evidence for the importance of 2C oligomerization. J. Mol. Biol. 236, (1994).

36. Lu, C. et al. Hexamers of the type II secretion ATPase GspE from Vibrio cholerae with Increased ATPase activity. Structure 21, (2013).

37. Monroe, N., Han, H., Shen, P. S., Sundquist, W. I. & Hill, C. P. Structural basis of protein translocation by the Vps4-Vta1 AAA ATPase. Elife 6, (2017).

38. Shi, H., Rampello, A. J. & Glynn, S. E. Engineered AAA+ proteases reveal principles of proteolysis at the mitochondrial inner membrane. Nat. Commun. 7, (2016).

39. Zaccai, N. R. et al. A de novo peptide hexamer with a mutable channel. Nat. Chem. Biol. 7, (2011).

40. Echeverri, A. C. & Dasgupta, A. Amino terminal regions of poliovirus 2C protein mediate membrane binding. Virology 208, (1995).

41. Bonafoux, D. et al. Fragment-Based Discovery of Dual JC Virus and BK Virus Helicase Inhibitors. J. Med. Chem. 59, (2016).

42. Tang, Q. et al. Identification of dibucaine derivatives as novel potent enterovirus 2C helicase inhibitors: In vitro, in vivo, and combination therapy study. Eur. J. Med. Chem. 202, (2020).

43. Bochkov, Y. A. et al. Molecular modeling, organ culture and reverse genetics for a newly identified human rhinovirus C. Nat. Med. 17, (2011).

44. Banerjee, S. et al. 2.3 Å resolution cryo-EM structure of human p97 and mechanism of allosteric inhibition. Science (80-.). 351, (2016).

45. Su, M. et al. Mechanism of Vps4 hexamer function revealed by cryo-EM. Sci. Adv. 3, (2017).

46. Zhao, M. & Brunger, A. T. Recent Advances in Deciphering the Structure and Molecular Mechanism of the AAA + ATPase N-Ethylmaleimide-Sensitive Factor (NSF). Journal of Molecular Biology vol. 428 (2016).

47. Lantez, V. et al. Comparative production analysis of three phlebovirus nucleoproteins under denaturing or non-denaturing conditions for crystallographic studies. PLoS Negl. Trop. Dis. 5, (2011).

48. Kabsch, W. et al. XDS. Acta Crystallogr. Sect. D Biol. Crystallogr. 66, (2010).

49. Emsley, P. & Cowtan, K. Coot: Model-building tools for molecular graphics. Acta Crystallogr. Sect. D Biol. Crystallogr. 60, (2004).

50. Afonine, P. V. et al. Real-space refinement in PHENIX for cryo-EM and crystallography. Acta Crystallogr. Sect. D Struct. Biol. 74, (2018).

51. Pettersen, E. F. et al. UCSF Chimera - A visualization system for exploratory research and analysis. J. Comput. Chem. 25, (2004).

52. Wessels, E. et al. Effects of Picornavirus 3A Proteins on Protein Transport and GBF1-Dependent COP-I Recruitment. J. Virol. 80, (2006).

53. Lanke, K. H. W. et al. GBF1, a Guanine Nucleotide Exchange Factor for Arf, Is Crucial for Coxsackievirus B3 RNA Replication. J. Virol. 83, (2009).

54. Reed, L. J. & Muench, H. A simple method of estimating fifty per cent endpoints. Am. J. Epidemiol. 27, (1938).

55. Zivanov, J. et al. New tools for automated high-resolution cryo-EM structure determination in RELION-3. Elife 7, (2018).

56. Zheng, S. Q. et al. MotionCor2: Anisotropic correction of beam-induced motion for improved cryo-electron microscopy. Nature Methods vol. 14 (2017).

57. Zhang, K. Gctf: Real-time CTF determination and correction. J. Struct. Biol. 193, (2016).

58. Tan, Y. Z. et al. Addressing preferred specimen orientation in single-particle cryo-EM through tilting. Nat. Methods 14, 793–796 (2017).

59. Su, M. goCTF: Geometrically optimized CTF determination for single-particle cryo-EM. J. Struct. Biol. 205, (2019).

60. Goddard, T. D. et al. UCSF ChimeraX: Meeting modern challenges in visualization and analysis. Protein Sci. 27, (2018).

