## Supplementary information for "Fluoxetine targets an allosteric site in the enterovirus 2C AAA+ ATPase and stabilizes the hexameric complex"

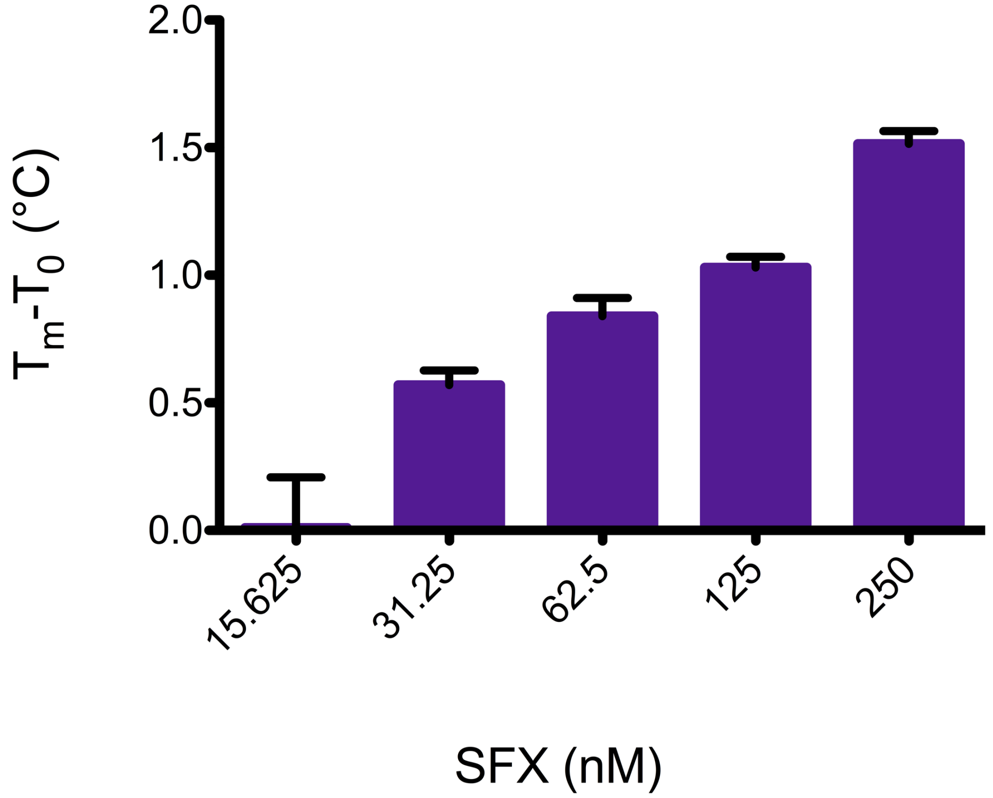


**Figure S1. (S)-Fluoxetine binds to the nonstructural protein 2C *in vitro*.** The binding of SFX to CVB3 Δ117-2C was assessed by thermal shift assay. The binding of SFX to Δ117-2C is represented by an increase in melting temperature, which indicates the thermal stabilization of the protein.

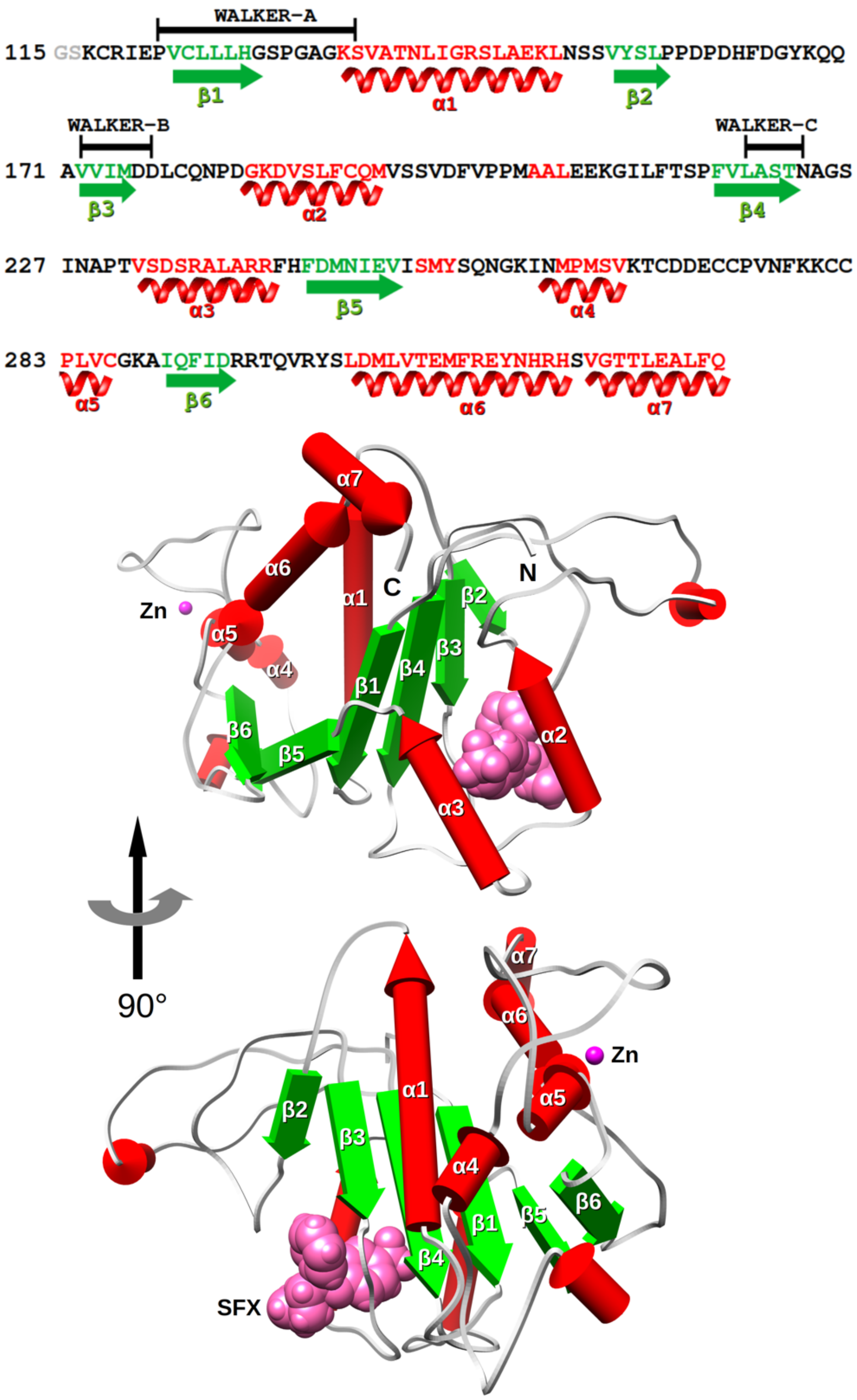


**Figure S2. Annotated sequence and structure of CVB3 Δ117-2C.** The fold of CVB3 2C in complex with *S*-fluoxetine (shown in pink color) is constituted by a Rossman fold core domain prolonged by a sub-4-Cys zinc finger domain and a C-terminal long helix. The β-sheet and α-helices respectively in green and red color are named accordingly to their position in the structure as depicted in the upper panel of the figure.

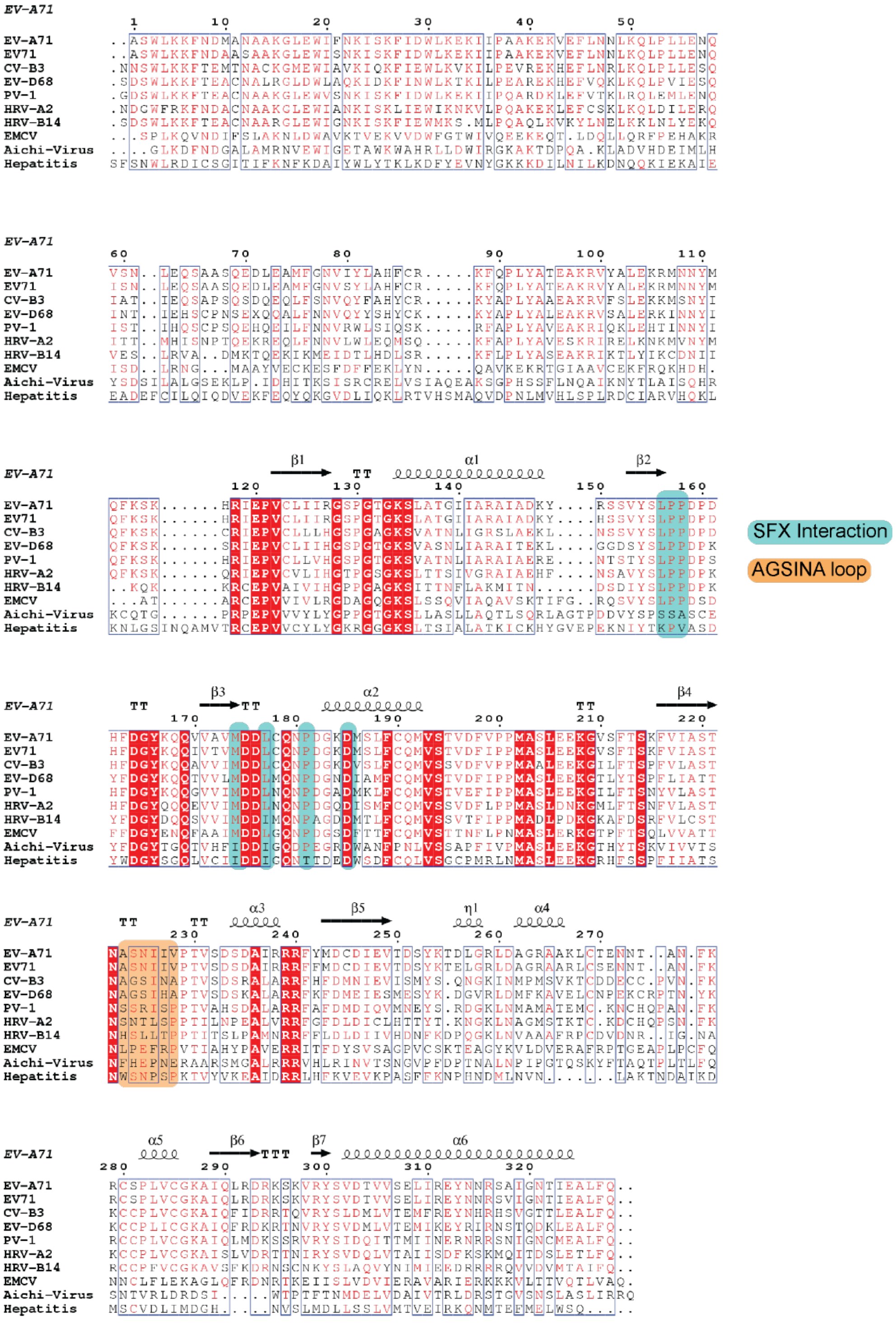


**Figure S1. Multiple sequence alignment of 2C proteins of different picornaviruses.** The sequence alignment of EV-A71 (BrCr), CV-B3 (strain Nancy), PV (strain Sabin), EV-D68 (strain Fermon), HRV-A2, HRV-B14, Encephalomyocarditis virus (EMCV), Aichi virus, and Hepatitis A virus was performed with ClustalOMEGA[1]. The alignment was subjected to the ESPript 3.0 server[2]. Conserved residues are highlighted in red with white letters. Highly conserved residues are highlighted in red letters. Secondary structural elements are shown on top of the alignment and are based on the EV-A71 crystal structure (PDB: 5GRB). Green Boxes indicate interaction residues with (*S*)-fluoxetine. The orange box indicated the AGSINA loops at position 224-229 in which SFX resistance mutations occur.

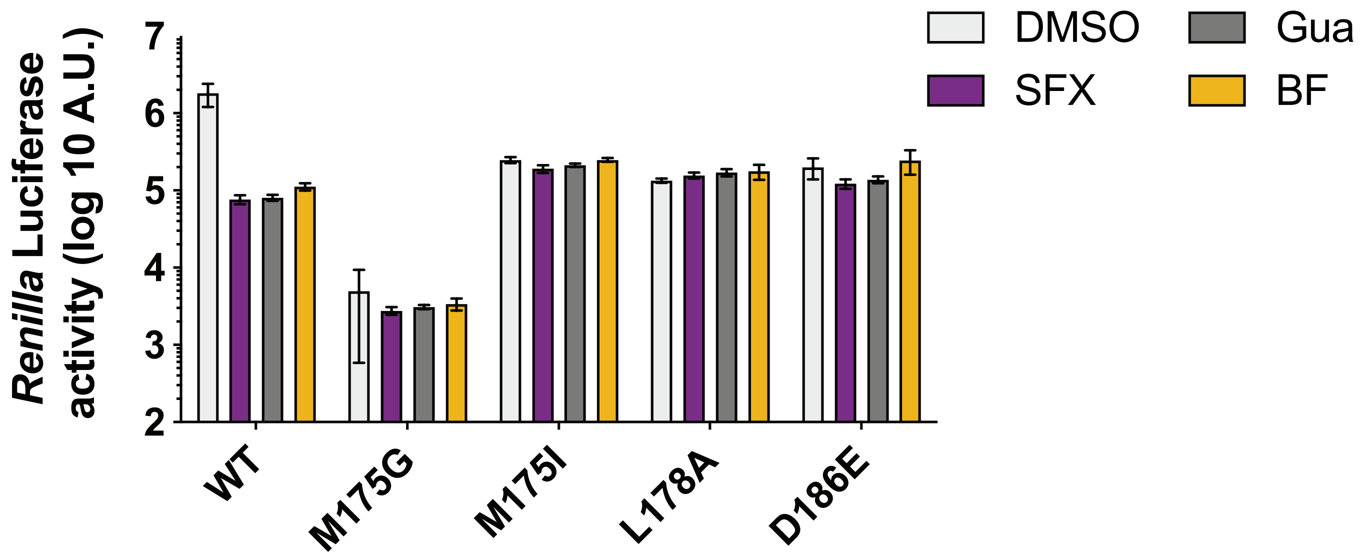


**Figure S4. Introduction of less stringent mutations in CV-B3 2C cannot rescue CV-B3 replication.** (A) Less stringent mutations were introduced into a recombinant CV-B3 virus encoding a Renilla luciferase reporter gene (Rluc-CV-B3) upstream of the capsid coding region. Infectious RNA was transfected into cells and *Renilla* luciferase was used as a sensitive and quantitative read-out for virus replication.

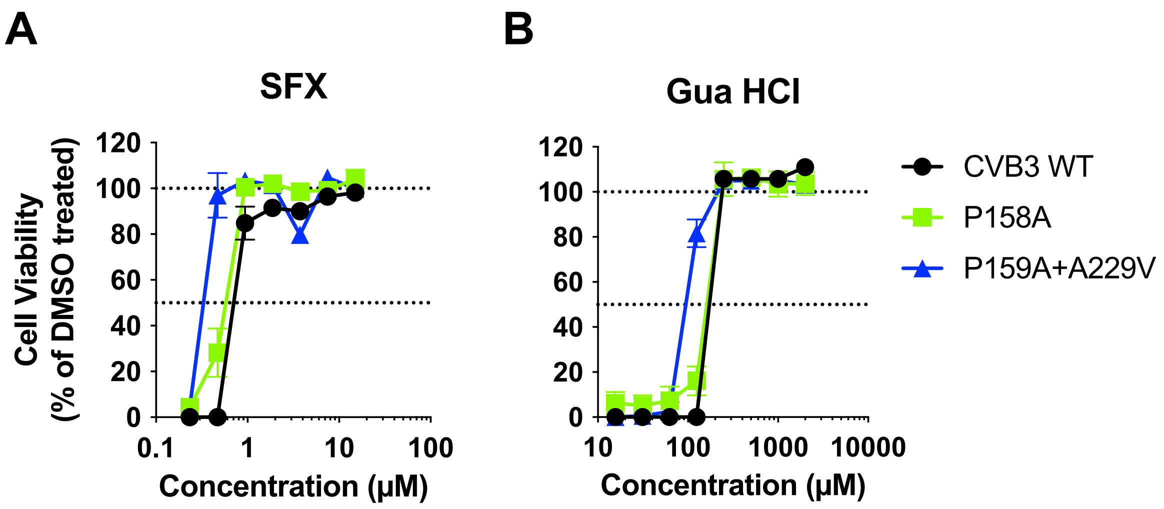


**Figure S5. Resistance profile of viable 2C mutations of CV-B3.** Viruses with several 2C mutations were tested for their sensitivity against (A) SFX or (B) GuaHCl in a multicycle replication assay. The experimental data displayed represent one out of three independent experiments which were performed in biological triplicates.

**A**

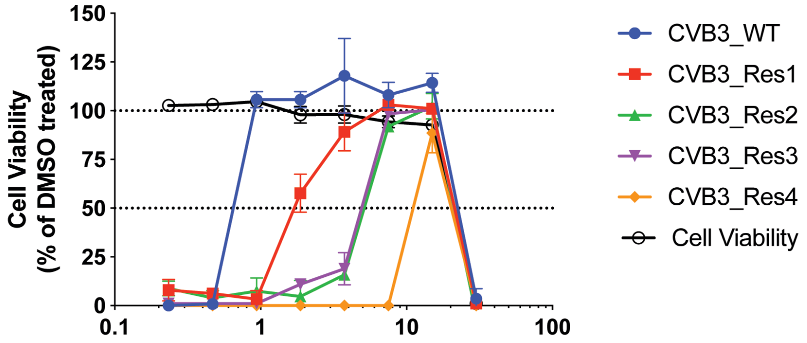


**B**

Resistant Virus Genotypes

| CVB3 | | Genotype 2C | SFX (µM) | Fold |
|---|---|---|---|---|
| | CVB3-WT | WT | 0,62 ± 0,06 | 1 |
| | CVB3 SFX-Res1 | I227V | 2,31 ± 0,16 | 4 |
| | CVB3 SFX-Res2 | I227V, A229V | 4,90 ± 0,25 | 8 |
| | CVB3 SFX-Res3 | I227V, A229V | 4,94 ± 0,05 | 8 |
| | CVB3 SFX-Res4 | A224V, I227V, A229V | 8,67 ± 0,68 | 14 |

**Figure S6. Raising SFX-resistant CVB3 viruses.** CV-B3 viruses resistant to S-fluoxetine (SFX) were raised in a multistep protocol as described previously (Bauer et al 2020). (A) Multicycle viral replication assay was performed to determine SFX sensitivity of CV-B3 viruses resistant to SFX. Therefore, HeLa R19 cells were treated with serial dilutions of SFX and infected with MOI of 0.001 After 3 days, the cells' viability was determined using an MTS assay. (B) Genotypes of 2C of the raised resistant CV-B3 viruses are shown

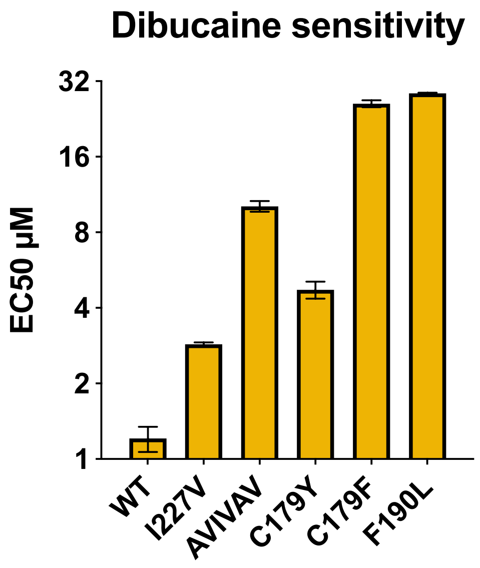


**Figure S7. 2C mutations providing resistance to SFX are cross-resistance to dibucaine.** Rluc CV-B3 reporter viruses containing previously identified mutations in the nonstructural protein 2C conferring resistance to several identified 2C inhibitors were used in a single cycle assay. HelaR19 cells were infected with a MOI 0.1 of Rluc-CV-B3 WT, the I227V mutant, the triple mutant A224V-I227V-A229V (designated AVIVAV), the C179Y of C179F mutant and the F190L mutant. One hour after infection, the cells were treated with a serial dilution of dibucaine. The 50% effective concentration $EC_{50}$ values displayed are calculated from three independent experiments which were performed in biological triplicates.

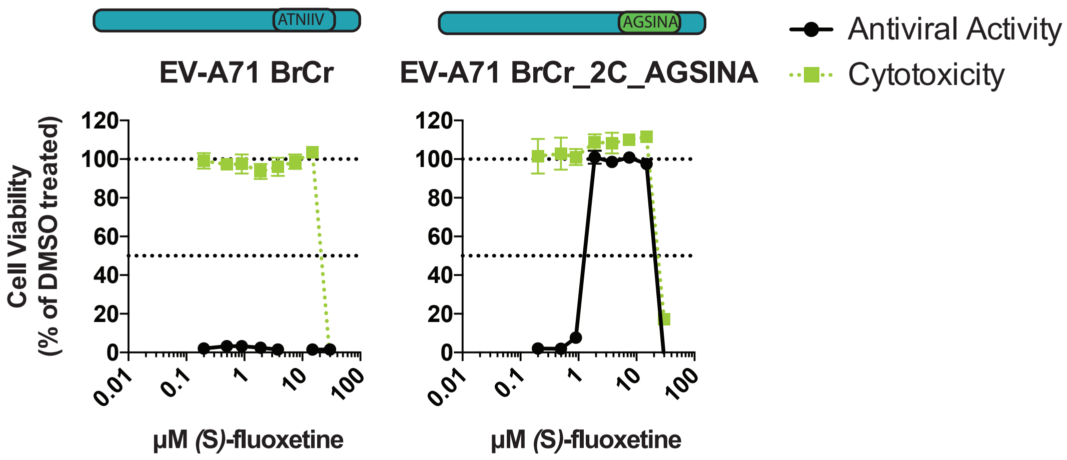


**Figure S8. Introduction of 224-AGSINA-229 into the infectious clone of EV-A71 results in SFX sensitivity.** The 224-AGSINA-229 loop was introduced into the 2C protein of the SFX-insensitive EV-A71 BrCr strain with reverse genetics. The obtained virus was used to determine the SFX-sensitivity in a multicycle assay. In parallel, the cytotoxicity of SFX was determined with a cell viability assay. The experimental data display represents one out of three independent experiments which were performed in biological triplicates.

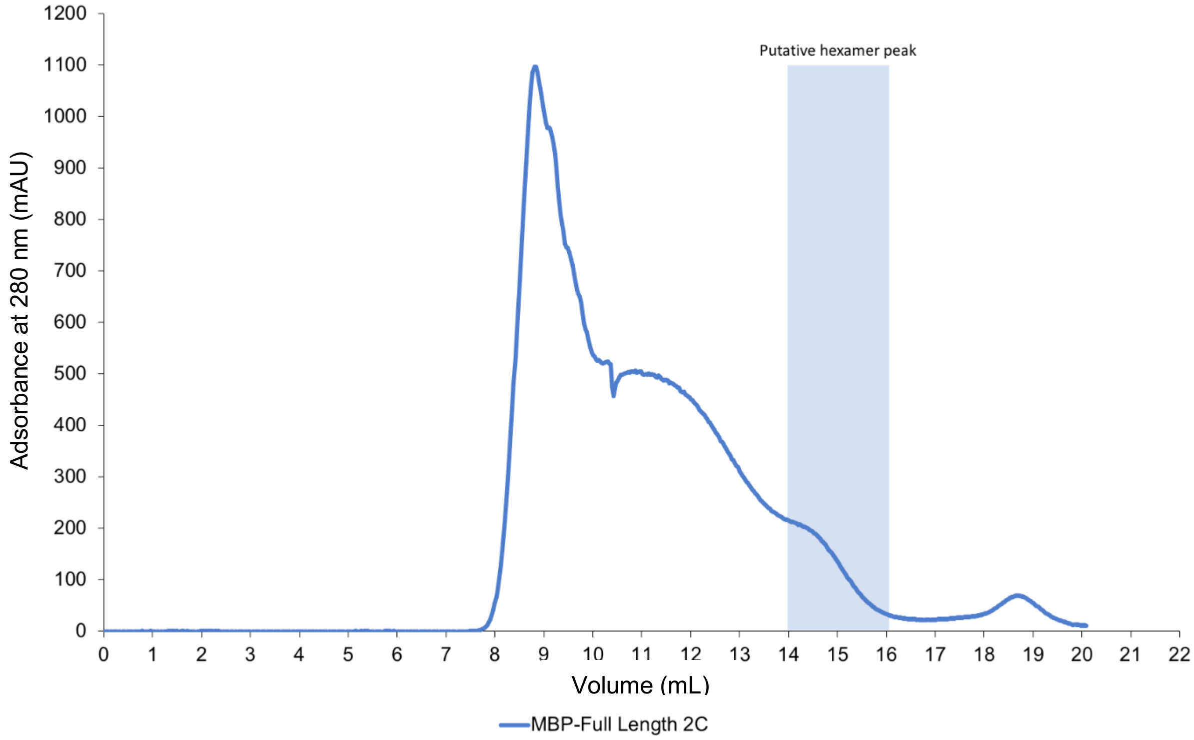


**Figure S9: Purification of MBP-tagged full-length CV-B2 2C protein.** Size Exclusion Chromatogram for the MBP-tagged full length 2C protein. Elution of the putative hexamer is expected between 14 and 16 mL with peak indicated.

6xhis

$H_3N^+$-MGSSHHHHHHMKIEEGKLVIWINGDKGYNGLAEVGKKFEKDTGIKVTVE

Maltose-binding protein

HPDKLEEKFPQVAATGDGPDIIFWAHDRFGGYAQSGLLAEITPDKAFQD

KLYPFTWDAVRYNGKLIAYPIAVEALSLIYNKDLLPNPPKTWEEIPALDKEL

KAKGKSALMFNLQEPYFTWPLIAADGGYAFKYENGKYDIKDVGVDNAG

AKAGLTFLVDLIKNKHMNADTDYSIAEAAFNKGETAMTINGPWAWSNIDT

SKVNYGVTVLPTFKGQPSKPFVGVLSAGINAASPNKELAKEFLENYLLTD

EGLEAVNKDKPLGAVALKSYEEELAKDPRIAATMENAQKGEIMPNIPQM

Linker HRV-3C

SAFWYAVRTAVINAASGRQTVDEALKDAQTGGGGSLEVLFQGPGGGG

Linker cc-hex Linker

SGGGGSGELKAIAQELKAIAKELKAIAWELKAIAQGAGGSGSYFQSNAS

CV-B3 2C (116-329)

KCRIEPVCLLLHGSPGAGKSVATNLIGRSLAEKLNSSVYSLPPDPDHFDG

YKQQAVVIMDDLCQNPDGKDVSLFCQMVSSVDFVPPMAALEEKGILFT

SPFVLASTNAGSINAPTVSDSRALARRFHFDMNIEVISMYSQNGKINMP

MSVKTCDDECCPVNFKKCCPLVCGKAIQFIDRRTQVRYSLDMLVTEMFR

EYNHRHSVGTTLEALFQ-$COO^-$

**Figure S10: Overview of the hexΔ116 CV-B3 2C protein expression construct.** Amino acid sequence of the hexΔ116-2C construct showing the positions of the 6xhis tag (red), MBP (green), HRV-3C protease cleavage site (blue), linker (grey), cc-hex-D24 sequence (purple) and residues 116-329 of CV-B3 2C (yellow).

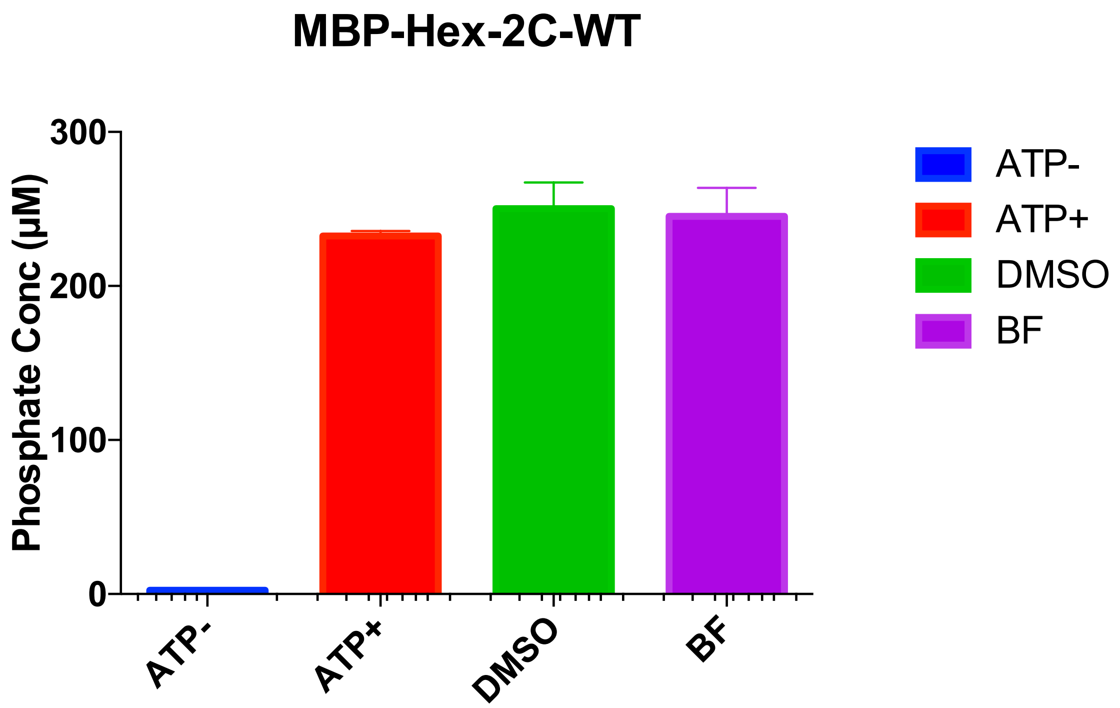


**Figure S11. Controls for the ATPase assay.** ATPase activity for the WT hexΔ116-2C with or without ATP added, and in the presence of DMSO or a non-2C targeting compound, BF.

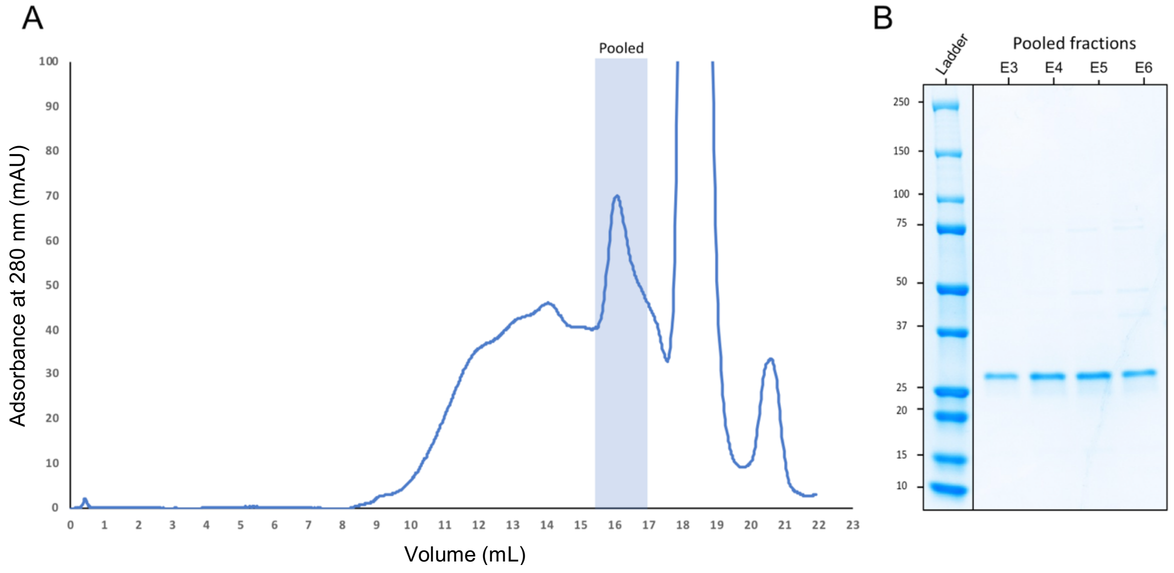


**Figure S12: Purification of the hexΔ116 CV-B3 2C protein following removal of the MBP tag.** A) Size Exclusion chromatogram of the hexΔ116-2C after removal of the MBP tag with 3C protease cleavage, with the pooled fractions indicated. B) Coomassie blue-stained SDS-PAGE analysis of pooled fractions.

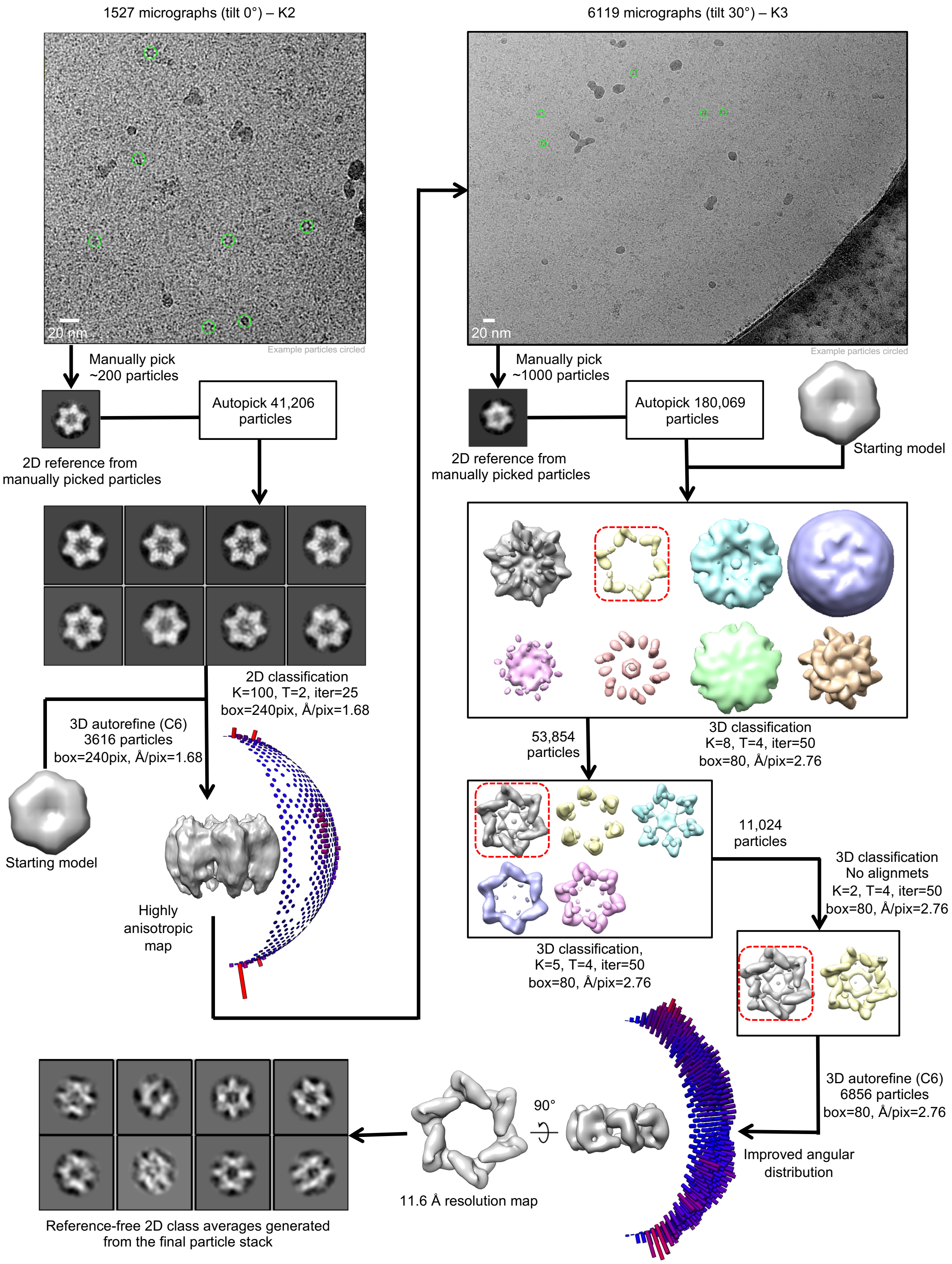


**Figure S13: Cryo-EM processing pipeline for hexΔ116 CV-B3 2C incubated with SFX.** Single-particle cryo-EM image processing workflow for hexΔ116-2C incubated with SFX (see methods for details).

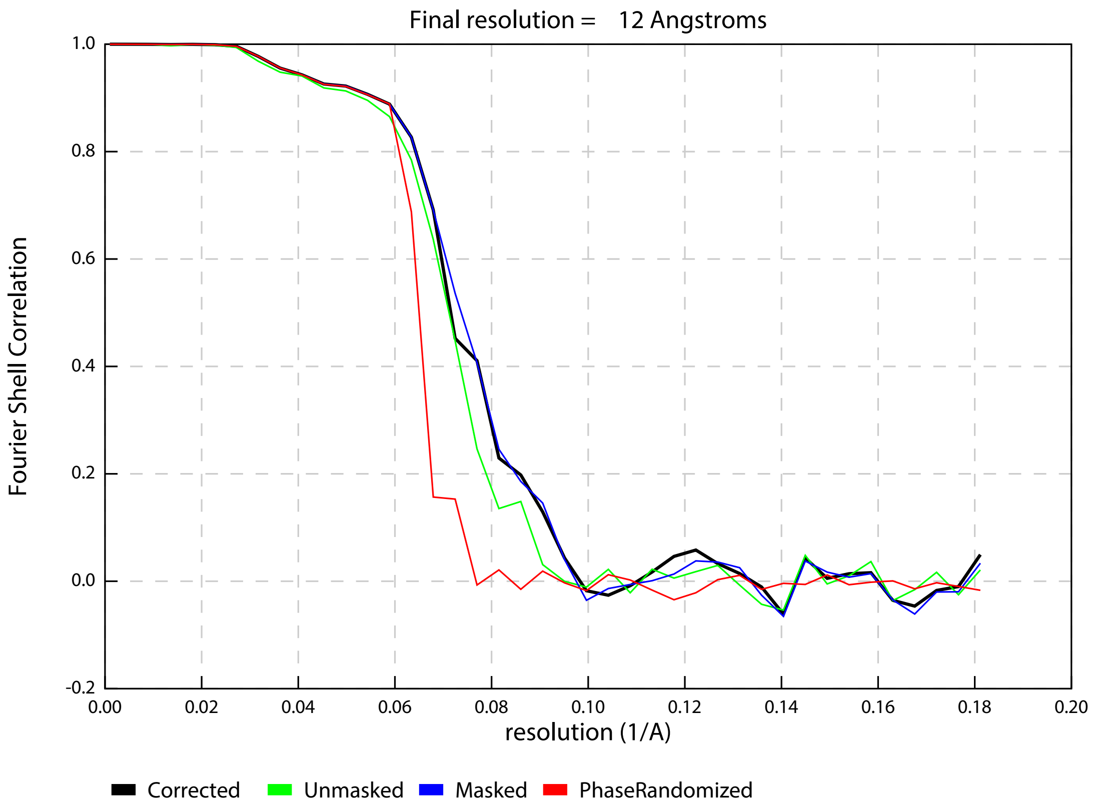


**Figure S14: Estimated resolution for the hexΔ116 CV-B3 2C cryo-EM reconstruction.** A) Gold-standard Fourier shell correlation (FSC) curve generated from the independent half maps contributing to the ~12 Å resolution density map.

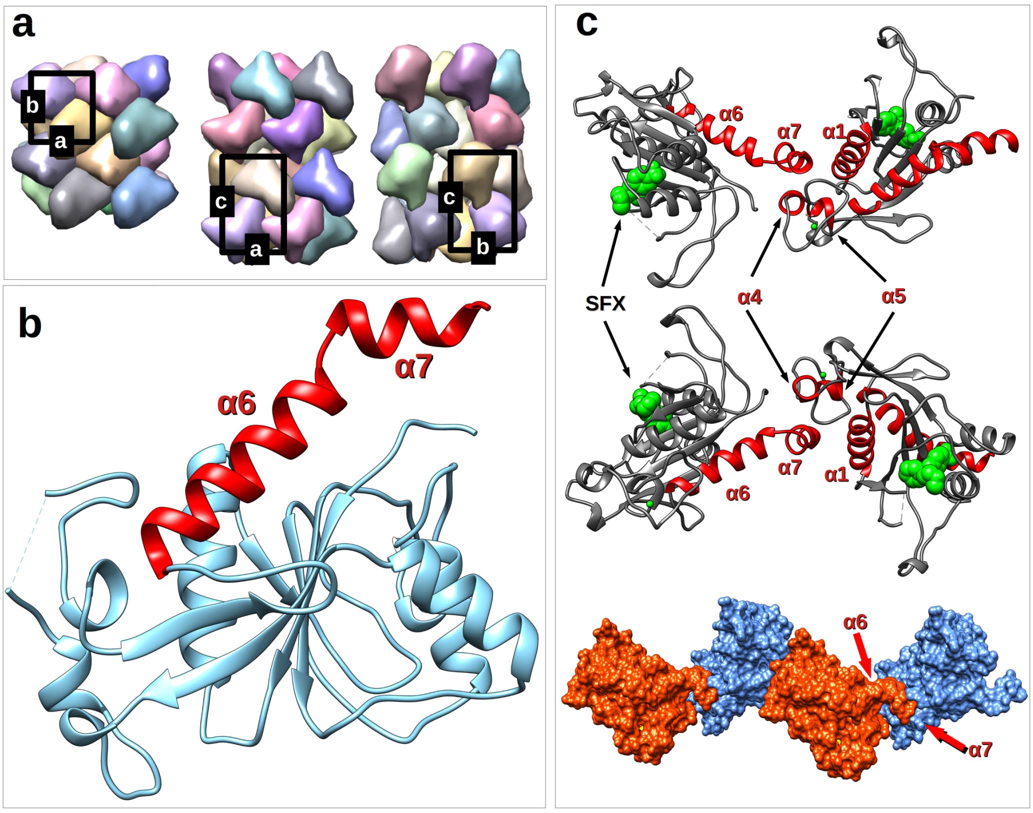


**Figure S15: Crystal packing of Δ117 CV-B3 2C in complex with SFX.** A) crystal packing of both structures reported here. B) The C-terminal of the CVB3 2C which probably acts as an oligomerization device. C) Crystal contact between symmetry related molecules showing the affinity of the C-terminal a7 helix towards the Zn binding site region of neighboring molecules within the crystal.

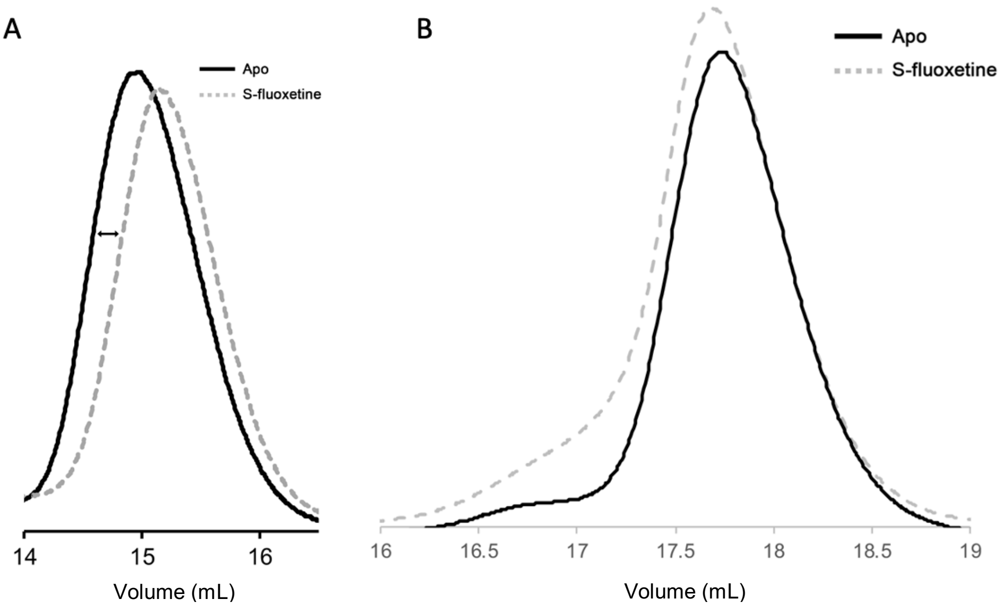


**Figure S16: Purification of hexΔ116 CV-B3 2C in the présence or absence of SFX.** Size Exclusion Chromatogram for the MBP-tagged hex116-2C protein in the presence or absence of 30 μM SFX. As shown in (A) for the monomeric MBP-tagged 116-2C protein.

**Table S1: Data collection and refinement statistics.** Statistics for the highest-resolution shell are shown in parentheses

| | 5S3A | 6T3W |
|---|---|---|
| **Wavelength** | 0,98 | 0,98 |
| **Resolution range** | 44.14 - 1.52 (1.574 - 1.52) | 39.68 - 1.82 (1.885 - 1.82) |
| **Space group** | P 21 21 21 | P 21 21 21 |
| **Unit cell** | 48.40 53.18 79.15 90 90 90 | 47.99 53.00 79.37 90 90 90 |
| **Total reflections** | 178322 (17165) | 230223 (17203) |
| **Unique reflections** | 32092 (3154) | 18771 (1846) |
| **Multiplicity** | 5.6 (5.4) | 12.3 (9.3) |
| **Completeness (%)** | 98.18 (94.87) | 99.97 (100.00) |
| **Mean I/sigma(I)** | 13.69 (1.89) | 21.73 (2.16) |
| **Wilson B-factor** | 16.39 | 29.35 |
| **R-merge** | 0.07018 (0.8355) | 0.06223 (0.8174) |
| **R-meas** | 0.07759 (0.9264) | 0.06496 (0.8656) |
| **R-pim** | 0.0326 (0.3955) | 0.01838 (0.2818) |
| **CC1/2** | 0.999 (0.646) | 1 (0.798) |
| **CC*** | 1 (0.886) | 1 (0.942) |
| **Reflections used in refinement** | 31556 (2994) | 18768 (1846) |
| **Reflections used for R-free** | 1889 (174) | 1878 (185) |
| **R-work** | 0.1853 (0.2538) | 0.1984 (0.2740) |
| **R-free** | 0.1995 (0.2579) | 0.2128 (0.3030) |
| **CC(work)** | 0.959 (0.800) | 0.954 (0.789) |
| **CC(free)** | 0.952 (0.798) | 0.945 (0.730) |
| **Number of non-hydrogen atoms** | 1873 | 1739 |
| **macromolecules** | 1623 | 1620 |
| **ligands** | 24 | 45 |
| **solvent** | 226 | 74 |
| **Protein residues** | 210 | 208 |
| **RMS(bonds)** | 0.007 | 0.013 |
| **RMS(angles)** | 1.09 | 1.63 |
| **Ramachandran favored (%)** | 98.36 | 98.53 |
| **Ramachandran allowed (%)** | 1.64 | 1.47 |

| | | |
|---|---|---|
| **Ramachandran outliers (%)** | 0.00 | 0.00 |
| **Rotamer outliers (%)** | 0.00 | 0.00 |
| **Clashscore** | 6.09 | 5.15 |
| **Average B-factor** | 21.15 | 36.76 |
| **macromolecules** | 19.69 | 35.99 |
| **ligands** | 25.48 | 50.67 |
| **solvent** | 31.11 | 45.10 |

**Table S2. Mutations in SFX coordinating residues does not result in replicating CV-B3.** Several 2C mutations were introduced into an infectious clone of CV-B3. Viral RNA was transcribed *in vitro* and transfected into HeLaR19 or BGM cells. + indicates full CPE observed, +/- indicates partial CPE, - indicates no CPE observed.

| | CPE<br>Sequencing result | | | | | | | | | | | | | | | | | |
|---|---|---|---|---|---|---|---|---|---|---|---|---|---|---|---|---|---|---|
| | HeLaR19 | | | | | | | | | BGM | | | | | | | | |
| | Passage 1 | | | Passage 2 | | | Passage 3 | | | Passage 1 | | | Passage 2 | | | Passage 3 | | |
| **Well** | 1 | 2 | 3 | 1 | 2 | 3 | 1 | 2 | 3 | 1 | 2 | 3 | 1 | 2 | 3 | 1 | 2 | 3 |
| **WT** | + | + | + | + | + | + | + | + | + | + | + | + | + | + | + | + | + | + |
| **L157A** | +<br>WT | - | - | +<br>WT | - | - | +<br>WT | - | - | - | - | - | - | - | - | - | - | - |
| **P158A** | +<br>P158A | +<br>P158A | +<br>P158A | +<br>P158A | +<br>P158A | +<br>P158A | +<br>P158A | +<br>P158A | +<br>P158A | +<br>P158A | +<br>P158A | +<br>P158A | +<br>P158A | +<br>P158A | +<br>P158A | +<br>P158A | +<br>P158A | +<br>P158A |
| **P159A** | - | +<br>P159A<br>A229V | - | - | +<br>P159A<br>A229V | - | - | +<br>P159A<br>A229V | +/-<br>P159A<br>A229V | - | - | - | - | - | - | - | - | - |
| **M175A** | - | - | - | - | - | - | - | - | - | - | - | - | - | - | - | - | - | - |
| **M175G** | - | - | - | - | - | - | - | - | - | - | - | - | - | - | - | - | - | - |
| **M175I** | - | - | - | - | - | - | - | - | - | - | - | - | - | - | - | - | - | - |
| **D176A** | - | - | - | - | - | - | - | - | - | - | +<br>WT | - | - | +<br>WT | - | - | +<br>WT | - |
| **D176N** | - | +<br>WT | - | - | +<br>WT | - | - | +<br>WT | - | - | - | - | - | - | - | - | - | - |
| **L178A** | - | - | - | - | - | - | - | - | - | - | - | - | - | - | - | - | - | - |
| **L178I** | - | - | - | - | - | - | - | - | - | - | - | - | - | - | - | - | - | - |
| **P182A** | - | - | - | - | - | - | - | - | - | - | +<br>WT | - | - | +<br>WT | - | - | +<br>WT | - |
| **D186A** | - | - | - | - | - | - | - | - | - | - | - | - | - | - | - | - | - | - |
| **D186N** | - | - | - | - | - | - | - | - | - | - | - | - | - | - | - | - | - | - |
| **D186E** | - | - | - | - | - | - | | | | - | - | - | - | - | - | - | - | - |

**Table S3 Primer list for 2C site directed mutagenesis.** Changed bases are indicated in lower cases.

| **2C mutations** | | **5'-3' sequence** |
|---|---|---|
| L157A | FWD | AGTGTACTCAgcaCCGCCAGACC |
| | REV | GAGCTGTTGAGTTTCTCAG |
| P158A | FWD | GTACTCACTAgcgCCAGACCCAG |
| | REV | ACTGAGCTGTTGAGTTTCTCAGC |
| P159A | FWD | CTCACTACCGgcaGACCCAGATC |
| | REV | TACACTGAGCTGTTGAGTTTCTC |
| M175A | FWD | CGTGGTGATTgcgGACGATCTATGC |
| | REV | GCCTGCTGTTTGTATCCG |
| M175G | FWD | CGTGGTGATTgggGACGATCTATGC |
| | REV | GCCTGCTGTTTGTATCCG |
| M175I | FWD | CGTGGTGATTattGACGATCTATGC |
| | REV | GCCTGCTGTTTGTATCCG |
| D176A | FWD | GGTGATTATGgccGATCTATGCC |
| | REV | ACGGCCTGCTGTTTGTAT |
| D176N | FWD | GGTGATTATGaacGATCTATGC |
| | REV | ACGGCCTGCTGTTTGTAT |
| L178A | FWD | TATGGACGATgcaTGCCAGAATCCTGATG |
| | REV | ATCACCACGGCCTGCTGT |
| L178I | FWD | TATGGACGATattTGCCAGAATCCTGATGG |
| | REV | ATCACCACGGCCTGCTGT |
| P182A | FWD | ATGCCAGAATgccGATGGGAAAG |
| | REV | AGATCGTCCATAATCACC |
| D186A | FWD | TGATGGGAAAgccGTCTCCTTGT |
| | REV | GGATTCTGGCATAGATCG |
| D186E | FWD | TGATGGGAAAgagGTCTCCTTGT |
| | REV | GGATTCTGGCATAGATCG |
| D186N | FWD | TGATGGGAAAaacGTCTCCTTGT |
| | REV | GGATTCTGGCATAGATCGTC |
| A229V | FWD | ATCTATTAATgttCCAACCGTGTCAG |
| | REV | CCTGCATTGGTCGATGCC |

**Table S4: Cryo-EM data collection and image processing**

| | Hex116-2C + SFX | |
|---|---|---|
| **Data Collection** | | |
| Camera | K2 | K3 |
| Detector mode | Counting | Super resolution |
| Movies collected | 1527 | 6119 |
| Magnification | 165 000 | 64 000 |
| Voltage (kV) | 300 | 300 |
| Stage tilt (°) | 0 | 30 |
| Electron exposure (e-/Å$^2$) | 50 | 54 |
| Defocus range (μm) | 1.5-2.5 | 2-4 |
| Pixel size (Å) | 0.84 | 0.69 |
| **Image processing** | | |
| Pixel size (Å) | 1.68 | 2.76 |
| Symmetry imposed | C6 | C6 |
| Initial particle images (no.) | 41 206 | 180 069 |
| Final particle images (no.) | 3616 | 6856 |
| Map resolution (Å) | N/A | 12 |
| FSC threshold | N/A | 0.143 |

### Supplementary References


1. Sievers, F. *et al.* Fast, scalable generation of high-quality protein multiple sequence alignments using Clustal Omega. *Mol. Syst. Biol.* **7**, (2011).
2. Robert, X. & Gouet, P. Deciphering key features in protein structures with the new ENDscript server. *Nucleic Acids Res.* **42**, (2014).
